# Daily light-induced transcription in visual cortex neurons drives downward Firing Rate Homeostasis and stabilizes sensory processing

**DOI:** 10.1101/2024.05.05.592565

**Authors:** Dahlia Kushinsky, Emmanouil Tsivourakis, Daniella Apelblat, Ori Roethler, Mor Breger-Mikulincer, Katayun Cohen-Kashi Malina, Ivo Spiegel

**Affiliations:** Department of Brain Sciences, Weizmann Institute of Science, Rehovot, Israel; Department of Molecular Neuroscience, Weizmann Institute of Science, Rehovot, Israel

**Keywords:** Experience-induced transcription, Visual cortex, Firing Rate Homeostasis, Synaptic plasticity, Inhibition, E/I-ratio, Npas4, Visual processing

## Abstract

Balancing plasticity and stability in neural circuits is essential for an animal’s ability to learn from its environment while preserving the proper processing and perception of sensory information. However, unlike the mechanisms that drive plasticity in neural circuits, the activity-induced molecular mechanisms that convey functional stability remain poorly understood. Focusing on the visual cortex of adult mice and combining transcriptomics, electrophysiology and 2-photon imaging, we find that the daily appearance of light induces in excitatory neurons a large gene program along with rapid and transient shifts in the ratio of excitation and inhibition (E/I-ratio) and ongoing neural activity. Furthermore, we find that the light-induced transcription factor NPAS4 drives these daily normalizations of E/I-ratio and neural activity rates and that it stabilizes the neurons’ response properties. These findings indicate that daily sensory-induced transcription normalizes E/I-ratio and drives downward Firing Rate Homeostasis to maintain proper sensory processing and perception.

## Introduction

Balancing plasticity and stability in neural circuits is crucial for an animal’s survival: neural circuits have to be plastic such that animals can adapt to and learn from their sensory experiences but they also have to remain functionally stable such that the proper processing and perception of sensory information is maintained and previously stored sensory information won’t be lost. However, while the molecular-cellular mechanisms that convey plasticity to neural circuits - e.g. via Hebbian synaptic plasticity mechanisms ^1^ - have been studied extensively, the activity-dependent molecular mechanisms that convey functional stability to neural circuits are less understood and their function *in vivo* in intact neural circuits is largely unexplored.

Theoretical studies have revealed that if unchecked, Hebbian synaptic plasticity can lead to positive feedback loops of synaptic strengthening which might lead to run-away-excitation and to changes e.g. in the response properties (i.e. “tuning”) of neurons and to altered sensory perception ^2–7^. These theoretical studies proposed that at least three activity-induced cellular mechanisms counteract Hebbian plasticity mechanisms to maintain the activity rates of neural circuits in a homeostatic manner around a circuit-intrinsic setpoint and to thereby convey functional stability to neural circuits ^8–10:^ (1) scaling of excitatory synapses, (2) modulation of neuronal excitability, and (3) adjustment of the ratio of excitatory and inhibitory inputs (E/I-ratio) impinging onto a neuron. Indeed, each of these homeostatic plasticity mechanisms ^11,12^ as well as firing rate homeostasis (FRH) *per se* were subsequently observed in various experimental settings including in dissociated cultured neurons ^13^ and in intact neural circuits *in vivo* ^14,15^. These experimental studies also revealed that the time course of homeostatic plasticity mechanisms is relatively slow compared to Hebbian mechanisms as they usually start manifesting only several hours after the initial stimulus. However, whether FRH actually occurs *in vivo* in response to naturally occuring stimuli during an animal’s everyday life remains unclear since FRH was demonstrated *in vivo* so far only upon drastic experimental manipulations (e.g. upon monocular deprivation in the rodent visual cortex ^14–18)^. Furthermore, it is still largely unclear whether and how specific aspects of neural circuit function (e.g. the processing, perception and/or storage of sensory information) are controlled by FRH. These gaps-in-knowledge are largely due to our poor understanding of the activity-induced molecular mechanisms that control FRH in the intact brain: while multiple genes and molecules were found to be required for cellular-synaptic homeostatic plasticity mechanisms in cultured neurons ^19–30^, the expression and/or activity of many of these genes and molecules are not directly regulated by neural activity *in vivo* and these non-activity-regulated genes and molecules are thus poor candidates for controlling FRH in an activity-dependent manner *in vivo* (they might, however, control circuit-intrinsic setpoints of FRH - see ^31^). Furthermore, even among those genes and molecules whose expression and/or activity are regulated by neuronal activity *in vivo*, the synaptic scaffolding molecule SHANK3 is to our knowledge the only gene or protein for which it was tested directly whether it regulates FRH in single neurons *in vivo* ^32^. Thus, the identification of activity-induced molecular mechanisms that control FRH *in vivo* in an intact neural circuit is important within itself and might allow for establishing which aspects of the processing, perception and/or storage of (sensory) information in the brain are controlled by FRH.

Sensory experience and the subsequent increase in neural activity induce in neurons within minutes the transcription of a large gene program ^33–36^ which includes a small set of early-induced transcription factors (15-20 TFs, including e.g. FOS and NPAS4) that reach their peak RNA levels within one hour after stimulus onset and that are shared between most types of neurons. These ubiquitously-induced TFs then activate in each type of neuron the transcription of a large cell-type-specific set of late-induced effector genes (up to ∼300 genes per cell type) which modulate the cellular properties and synaptic connectivity of the cells in which they are expressed ^37–39^. Early studies, based e.g. on loss-of-function approaches in which the respective gene was removed (“knocked out”) in the whole animal from birth, gave rise to the idea that experience-induced genes enhance the plasticity of neural circuits since the lack of these genes was associated with deficits in Hebbian forms of synaptic plasticity (e.g. long-term plasticity or depression [LTP or LTD, respectively] ^40–42)^, disrupted the experience-dependent development of neural circuits ^43–46^, and led to deficits in various learning and memory paradigms ^42,47–49^. However, recent studies challenge this view and indicate that the short-term (i.e. acute) cellular function of many experience-induced genes lies in the activity-induced cell-type-specific regulation of E/I-ratio ^28,37,38,50–54^. Since these effects on E/I-ratio seem to follow a homeostatic logic - i.e. they are predicted to increase inhibition onto a circuit’s principal neurons to thereby reduce neural activity of these neurons in an activity-dependent manner - these findings suggest that activity-induced transcription might play a role in FRH. However, this idea remains untested and it is not known whether activity-induced transcription indeed down-regulates neural activity in a homeostatic manner *in vivo* in the brain and how this might affect e.g. the processing and/or storage of sensory information in the adult brain.

To address these knowledge gaps, here we focused on the visual cortex of adult mice and tested whether experience-induced gene expression controls FRH during an animal’s everyday life to thereby stabilize sensory processing in the adult brain. By taking advantage of the naturally occuring daily cycle of dark and light, we demonstrate that the daily appearance of light (but not other factors such as e.g. the passage of time) drives in visual cortex excitatory neurons (1) the rapid induction of a large gene program, (2) a rapid and transient shift in the E/I-ratio in these neurons towards excitation which occurs via the rapid light-induced increase of excitatory inputs onto these neurons and the slower increase of inhibitory inputs, and (3) a rapid and transient increase in ongoing activity of these neurons that is normalized towards the end of the day’s light phase. Furthermore, we demonstrate that these daily normalizations of E/I-ratio and neural activity rates are driven by the light-induced transcription factor NPAS4 and that NPAS4 stabilizes specific visual response properties in excitatory neurons. Thus, our study identifies - to our knowledge for the first time - an activity-induced molecular mechanism that drives downward FRH *in vivo* in response to a naturally occurring stimulus and that is necessary for maintaining proper sensory processing in the adult brain.

## Results

### The daily appearance of light induces a selective gene program in visual cortex neurons

Sensory experience induces in neurons in the visual cortex of adult mice a large and cell-type-specific gene program ^37,39^. For example, prolonged dark-housing followed by exposure to regular structured light induces cell-type-specific gene programs that include genes encoding for ubiquitous early-induced TFs (e.g. *Npas4*, *Fos*) that reach their peak RNA expression within one hour after the appearance of light and cell-type-specific sets of late-induced effector genes (e.g. *Bdnf*, *Igf1*) that reach their peak expression typically a few hours later (for many genes within three to eight hours). This suggests that similar light-induced gene expression might be observed every day in visual cortex neurons when daylight appears after the night’s darkness. To test this idea in a rigorous manner, we housed adult mice in a standard 12/12 hour dark-light cycle and sacrificed the animals on the day of the experiment at multiple time points, starting from the moment just before turning on the lights (i.e. Zeitgeber time 0 [ZT0]) and continuing throughout the light period and the ensuing dark period (Dark-Light-Dark [DLD] cohorts, 2 males and 2 females per time point; **Figure 1A**); we then extracted total RNA from the visual cortices of these mice and conducted RNA-seq (see Methods). To differentiate between the effects on gene expression that were caused by the appearance (and ensuing disappearance) of daylight and those that were driven by other factors during the same period of time (e.g. by circadian mechanisms) we included in these experiments parallel cohorts of mice that were of identical age and sex and were housed under identical conditions but in which the light was not turned on during the day of the experiment (i.e. Dark-Dark-Dark [DDD] cohorts; **Figure 1A**). Importantly - and consistent with previous studies ^55,56^ - we confirmed that the additional dark period did not alter the motor activity of DDD mice (i.e. running on treadmill; **Supplemental Figure 1A-C**) and that the arousal state (assessed by pupillometry [**Supplemental Figure 1D**] and whisking [**Supplemental Figure 1E**]) does not change during the light phase in DLD mice. Thus, potential differences in gene expression in the visual cortices of DLD- and DDD-mice are due to the presence or absence of light between ZT 0 and ZT 12 on the day of the experiment but not due to other factors.

**Figure 1.**
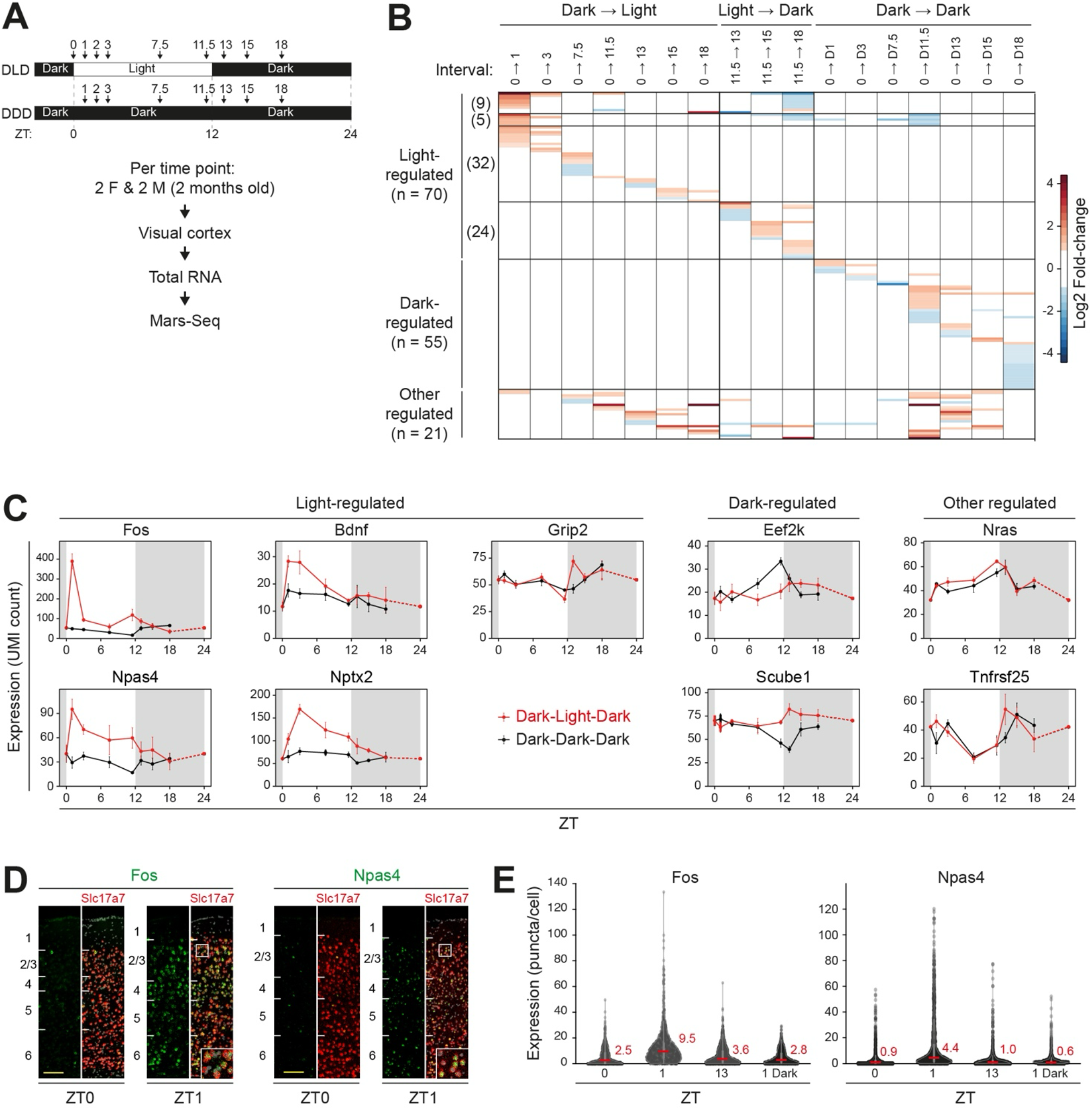
The daily appearance of light induces a selective gene program in visual cortex neurons. **A-C** RNA-seq on total RNA extracted from the visual cortices of adult mice identifies three classes of daily genes: 1. Light-regulated genes (i.e. genes that are regulated only by the transition from dark to light or light to dark [i.e. regulated only DLD cohorts]; n=70), 2. Dark-regulated genes (i.e. genes that are regulated only in mice that remained in the dark on the day of the experiments [DDD cohorts; n=55), 3. Other regulated genes (i.e. genes that are regulated in DLD and DDD cohorts; n=21). **A** Experimental strategy (ZT = Zeitgeber time; numbers & arrows indicate time point at which the animals were sacrificed; animals per time point = 2 males & 2 females per time point; DLD = Dark-Light-Dark cohort; DDD = Dark-Dark-Dark cohort). **B** Heatmap indicating the fold-changes of all the regulated genes at the respective time intervals across the day (white = no change). **C** Line plots of the expression of example genes for all three categories (error bars represent SEM in all panels). **D, E** RNAscope smFISH reveals that the expression of the early-induced TFs FOS and NPAS4 is induced by the daily appearance of light in excitatory neurons across all layers of the visual cortex. **D** Example images of coronal sections of the visual cortex with RNAscope smFISH for *Fos* or *Npas4* (green) and the excitatory marker *Slc17a7* (= VGlut1) at ZT0 (i.e. before light onset) or ZT1 (i.e. 1 hour after light onset). Numbers on the left indicate cortical layers, inlets are magnifications of boxed areas (scale-bar = 100 micron). **E** Quantification of the number of *Fos* and *Npas4* puncta in all visual cortex excitatory neurons at ZT0/1/13 in DLD mice or ZT1 in DDD mice (red line & number = median; see Supplemental Figure 1G for layer-specific quantification).

This transcriptomic approach led to the identification of 146 regulated genes (**Figure 1B,C**), 70 genes of which are regulated only by light (“light-regulated”, i.e. are regulated only in the visual cortices of DLD mice but not in DDD mice). These light-regulated genes include many genes that encode for well-known early-induced TFs (e.g. *Fos*, *Npas4*, *Egr1*/*2*/*4*) and late-induced effectors (e.g. *Bdnf*, *Nptx2*) (see also **Supplemental Table 1**) and that exhibit increases in their expression upon the appearance of light (i.e. 46 genes that change from ZT0 to ZT1/3/7.5/11.5/13/15/18) as well as some genes whose expression changes upon the transition from the light to the dark (24 genes that change from ZT11.5 to ZT13/15/18, e.g. *Grip2*) (**Figure 1B,C** and **Supplemental Data Table 1**). Consistent with previous findings on sensory-induced gene expression in the mouse visual cortex ^37,39,57^ and further indicating the validity of our transcriptomic data, our analyses also revealed that the RNA levels of many housekeeping genes do not change substantially across a single day (e.g. *ActB*, *Htt*, *Ubc*; **Supplemental Figure 1E**). As revealed by GO term analyses, the light-regulated genes are enriched for biological functions related to neurobiological processes and mechanisms related to neural plasticity, consistent with similar analyses of sensory-regulated gene programs in previous studies ^37,39^ (**Supplemental Figure 1F** and **Supplemental Data Table 1**). Interestingly, our transcriptomic analyses also identified a large set of genes (55 genes) whose RNA levels were regulated selectively in the visual cortices of DDD mice (“dark-regulated” genes; **Figure 1B,C**). However, unlike for the light-regulated genes, GO term analyses did not identify specific (neuro-)biological processes in which these dark-regulated genes might be involved. Finally, we identified 21 genes that are regulated both in DLD- and DDD-mice: as expected, this group included several genes previously found to be regulated in a circadian manner (e.g. *Tnfrsf25* ^58,59^).

Since we conducted our transcriptomic analyses in a non-cell-type-specific manner (i.e. bulk RNA-Seq on total RNA extracted from homogenized visual cortices), we compared our transcriptomic data to previously generated single-cell RNA-seq data of sensory-induced gene expression in the mouse visual cortex ^39^ to glean potential insight into the cell-type-specificity of the daily light-induced gene program (**Supplemental Data Table 1**). This revealed that, consistent with the large percentage of glutamatergic neurons in the cortex (∼85% of all cortical neurons), most of the light-regulated genes identified by us and this previous study are regulated in cortical excitatory neurons while only few of these genes seem to be regulated every day in other cell types (e.g. *Cyr61* and *Klf4* selectively in Astrocytes, *Kras* selectively in VIP INs). Since our transcriptomic data are particularly informative regarding visual cortex excitatory neurons, we next used RNAscope single molecule Fluorescent In Situ Hybridization (RNAscope smFISH) to validate that the genes encoding for the early-induced TFs FOS and NPAS4 are indeed light-induced in visual cortex excitatory neurons and to test whether this occurs in specific layers of the visual cortex. This revealed that these two TFs are induced every day in visual cortex excitatory neurons selectively upon the appearance of light (i.e. only in DLD mice but not in DDD mice; **Figure 1D,E**) across all cortical layers (**Supplemental Figure 1G**) and that the majority of excitatory neurons express these genes every day to some degree (i.e. 95.3% and 85.8% of all excitatory neurons respectively express *Fos* and *Npas4* at ZT1). When we conducted similar RNAscope smFISH experiments for *Igf1*, a gene that we did not identify in our transcriptomic analyses as regulated but that we and others have previously identified as a late-induced effector gene in a sparse subtype of GABAergic interneurons that express the vasoactive intestinal peptide (VIP INs ^28,37,39,57^), we found that also this gene is induced every day in a light-dependent manner (**Supplemental Figure 1H**). Thus, our bulk transcriptomic analyses likely underestimate the total number of daily (light-)regulated genes in the visual cortex since they lack the cell-type-specificity to detect all daily (light-induced) changes in gene expression in the various subtypes of visual cortex neurons. We therefore conclude that the appearance of daylight induces in visual cortex neurons every day a large light-dependent gene program.

### Daily light drives in visual cortex excitatory neurons a rapid increase in excitatory inputs and shift in E/I-ratio towards excitation which is normalized by a slow increase in inhibitory inputs

Gene expression programs determine the cellular properties and synaptic connectivity of neurons. Thus, since we found that the appearance of daylight induces a program of gene expression in visual cortex excitatory neurons, we next tested whether the daily appearance of light alters the cellular-synaptic properties of these neurons as well. For this, we used whole-cell patch clamp electrophysiology in acute visual cortex slices and focused our recordings on excitatory pyramidal (PYR) neurons in layer 2/3 (L2/3) of the visual cortex. As in our gene expression analyses (**Figure 1** and **Supplemental Figure 1**), we housed adult wild-type mice in a standard 12/12 dark-light cycle (i.e. DLD cohort), sacrificed the mice at ZT 0 (i.e. just before the daily appearance of light) or at multiple other time points throughout light phase of the day (ZT 1, 4, 8, 12), prepared acute visual cortex slices and then recorded from visually identified L2/3 PYR neurons; as before, we included in these experiments also DDD mice, i.e. mice that remained in the dark on the day of the experiment (**Figure 2A**).

**Figure 2.**
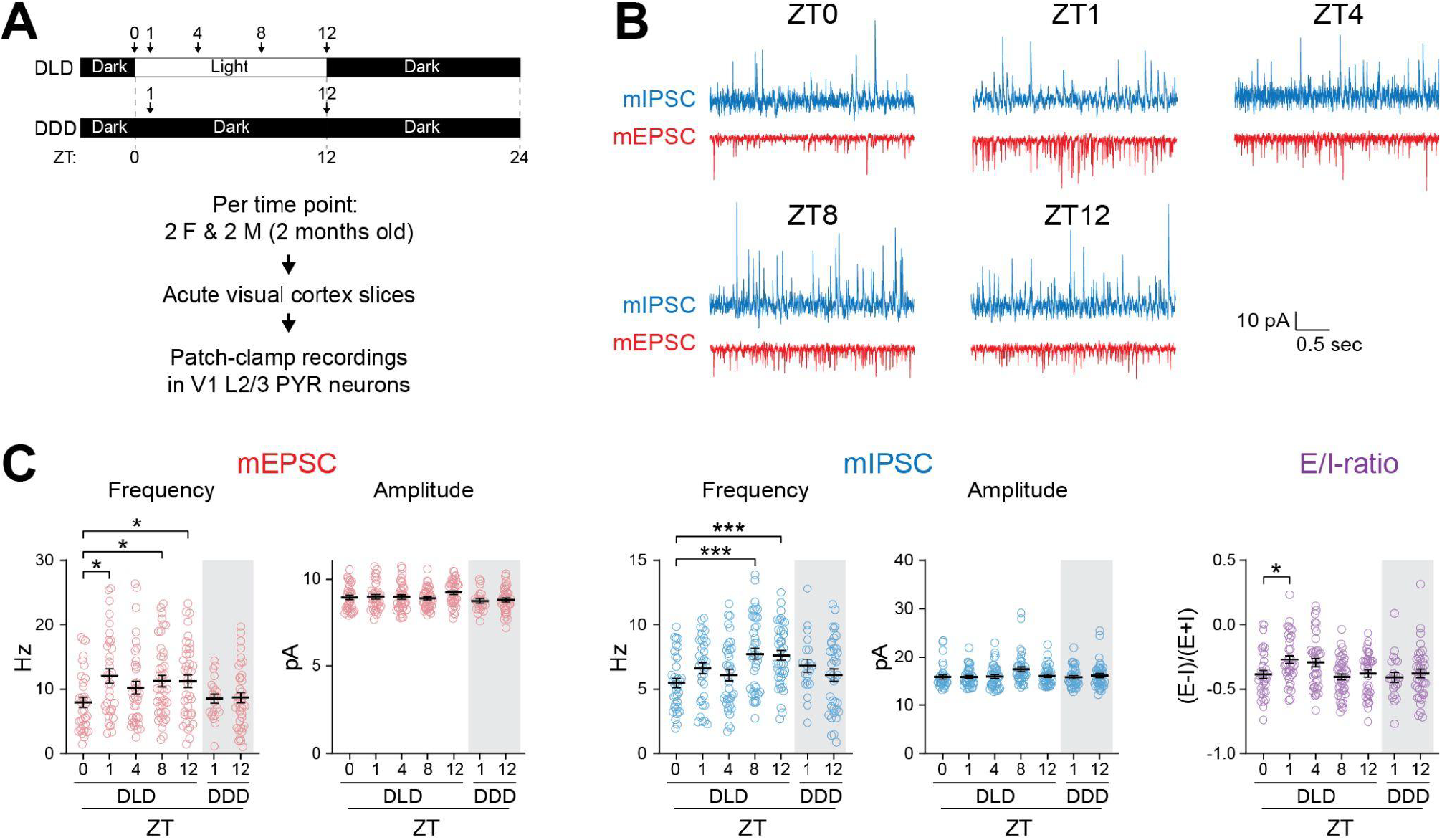
Daily light drives in visual cortex excitatory neurons a rapid increase in excitatory inputs and a shift in E/I-ratio towards excitation which is normalized by a slow increase in inhibitory inputs. **A** Experimental strategy for whole cell patch clamp recordings (ZT = Zeitgeber time; numbers & arrows indicate ZT at which the animals were sacrificed; n = 4 mice per ZT; DLD = Dark-Light-Dark cohort; DDD = Dark-Dark-Dark cohort). **B** Representative traces of miniature excitatory (red) and inhibitory (blue) postsynaptic currents (mEPSCs and mIPSCs, respectively) recorded from visual cortex L2/3 PYR neurons at different ZTs throughout the day. **C** Quantified data of frequencies and amplitudes of mIPSCs and mEPSCs and of E/I-ratio (For each time point, cells were recorded from 4 mice [4-9 cells/mouse]; Number of cells - DLD: ZT0 n= 37, ZT1 n= 39, ZT4 n= 37, ZT8 n= 43, ZT12 n= 40. DDD: ZT1 n= 21, ZT12 n= 38; Statistics Brown-Forsythe ANOVA test; post-hoc Dunnett’s T3 multiple comparisons test, * p<0.05, ** p<0.01, *** p<0.005). (Error bars represent SEM in all data panels; see **Supplemental Data Table 2** for descriptive statistics)

To identify daily changes in excitatory and inhibitory inputs onto L2/3 PYR neurons, we recorded miniature excitatory and inhibitory postsynaptic currents (mEPSCs and mIPSCs) in each neuron by switching the holding potential back-and-forth between −70 mV and 0 mV (**Figure 2B**), thereby allowing us to calculate the E/I-ratio for each cell in a precise manner (see Methods). These recordings revealed that the appearance of light leads to a rapid increase in mEPSC frequency (but not in amplitude) by ∼25% within one hour after light onset and that E/I-ratio shifts towards excitation by ∼40% at the same time since no concurrent significant change in mIPSCs occurs (**Figure 2C**). Importantly, no such changes were observed in DDD mice, demonstrating that the changes in mEPSC frequency and E/I-ratio observed in the DLD cohorts were driven by the appearance of light but not by the passage of time. Further supporting the notion that the changes in E/I-ratio observed in DLD mice are caused specifically by the appearance of light, we did not observe an effect of the time that passed between the preparation of the slices and the actual recordings of the cells on the E/I-ratio (**Supplemental Figure 2A**). Our analyses further revealed that once elevated, mEPSC frequency remained elevated for the remaining hours of the light phase. Similar to the light-induced increase in mEPSC frequency, also the frequency of the mIPSCs increased selectively in a light-dependent manner (i.e. mIPSCs did not change in DDD mice) - however, unlike the rapid increase in mEPSC frequency within one hour after lights on, the increase in mIPSC frequency was much slower and reached its peak only at ZT12 (increase by ∼30%). In turn, this slow increase in mIPSCs gradually shifted E/I-ratio back towards inhibition such that by ZT8, E/I-ratio returned to the same values as before the light onset. Notably, we did not observe any significant changes in the intrinsic electrophysiological properties of L2/3 PYR neurons (rheobase, resting membrane potential, input resistance, action potential (AP) amplitude; **Supplemental Figure 2B-F**) across the day. Thus, taken together, these experiments demonstrate that the daily appearance of light (but not the passage of time) leads in visual cortex L2/3 PYR neurons (1) to a rapid and selective increase in mEPSC frequency and to a concomitant rapid shift in E/I-ratio towards more excitation within one hour after the appearance of light, and (2) to a slower increase in mIPSC frequency which normalizes the initial light-induced shift in E/I-ratio after several hours, but (3) to no detectable changes in the excitability of these neurons.

### The daily appearance of light drives a rapid and transient increase in the activity of visual cortex neurons

The relative amount of excitation and inhibition impinging onto a neuron determines its propensity to fire action potentials (i.e. its activity rates) ^60–62^. Thus, having found that the daily appearance of light causes a rapid and transient shift in E/I ratio towards excitation in L2/3 PYR neurons *ex vivo* in acute visual cortex slices, we next asked whether the activity of these neurons changes in a corresponding light-dependent manner every day *in vivo* in the visual cortex of adult behaving mice. To address this question, we took a similar approach as in our transcriptomic and electrophysiological experiments and performed longitudinal 2-photon calcium imaging in L2/3 PYR neurons in the monocular zone of the primary visual cortex (mV1) of head-fixed awake behaving adult mice. For this, we expressed the genetically encoded calcium indicator (GECI) GCaMP7f selectively in excitatory neurons in mV1 of adult wild-type mice via a stereotactically injected AAV construct and imaged the GCaMP-expressing neurons in L2/3 of mV1 at ZT 0, ZT 2 and ZT 12 via 2-photon microscopy while monitoring the animals’ arousal state (via pupillometry and monitoring of facial movement) and while either presenting them with visual stimuli to assess the neurons’ response properties or while measuring ongoing activity when no visual stimulus is presented (**Figure 3A**). Notably, we restricted our imaging sessions to ZT 0, ZT 2 and ZT 12 since we observed at these time points the largest daily light-driven changes in E/I-ratio in our electrophysiology experiments and since we wanted to reduce the animals’ stress and let them rest as much as possible between the imaging sessions (the animals were returned to their home cages between imaging sessions). Importantly, we precisely aligned (“matched”) the imaged field-of-view in each imaging session which enabled us to track the same individual neurons across all imaging sessions and to compare their activity and response properties across all time points (see Methods). Data analyses were done with our established analyses pipelines ^28,63^, and we included also in these imaging experiments a separate cohort of mice that remained in the dark on the day of the experiment (DDD mice) such that we could determine whether potential changes in the neurons’ activity were caused by the daily appearance of light.

**Figure 3.**
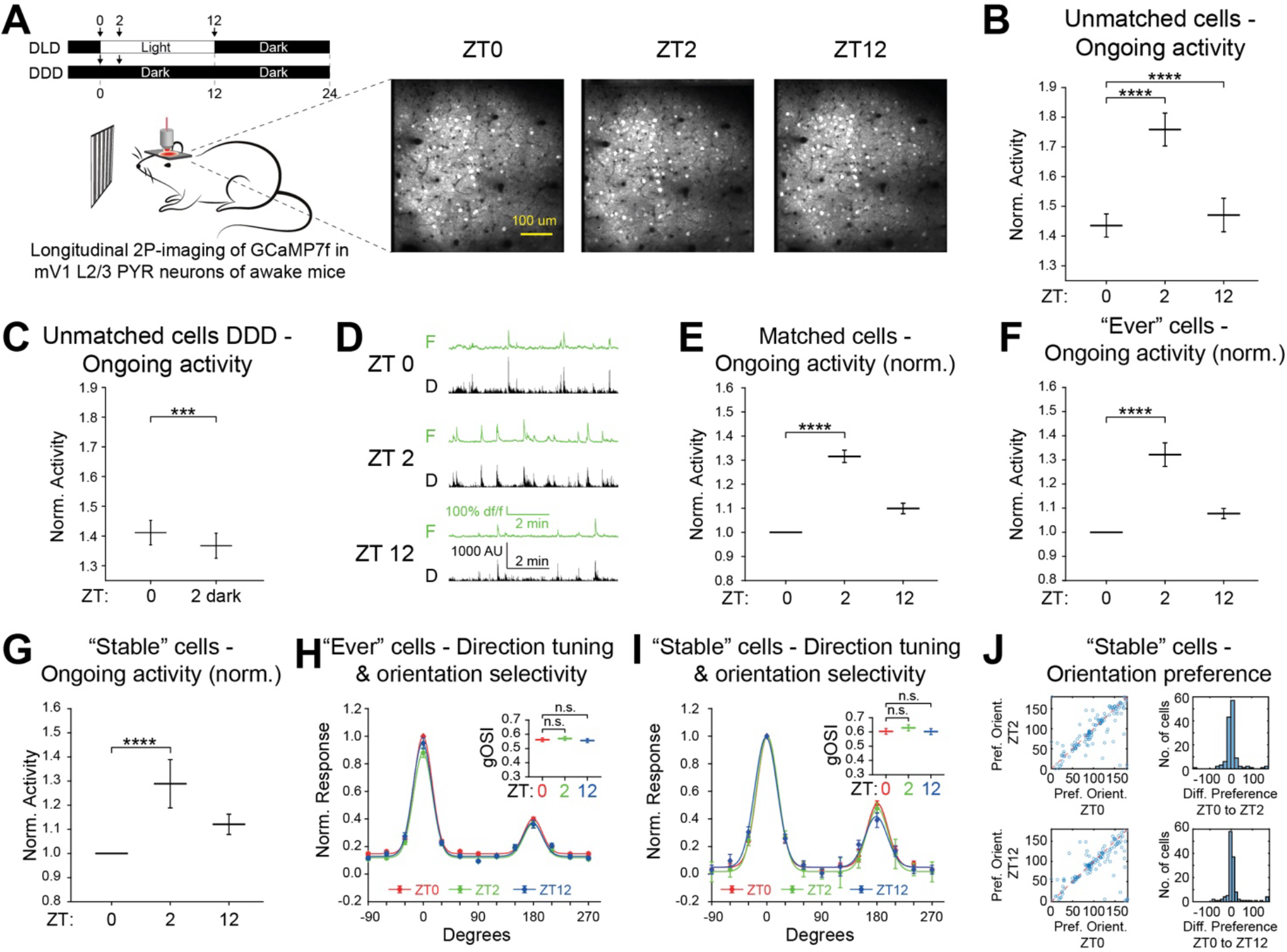
The daily appearance of light drives a rapid and transient increase in the ongoing activity of visual cortex neurons but does not alter tuning properties. **A (Left)** Experimental strategy for longitudinal in vivo 2-photon GCaMP-imaging of the same neurons across the day in the monocular zone of the primary visual cortex of awake, head-fixed mice (ZT = Zeitgeber time; numbers & arrows indicate ZT of recording sessions; mice were returned to their home cage between recordings; DLD = Dark-Light-Dark cohort; DDD = Dark-Dark-Dark cohort). **(Right)** Representative field of view (FOV) of a longitudinally recorded animal at ZT0, ZT2, and ZT12 (scale bar = 100 micron). **B** Daily changes in ongoing activity in all unmatched cells in DLD mice (cells recorded from 7 mice; ZT0: n = 7624 cells; ZT2: n = 7326 cells; ZT12: n = 7198 cells; Statistics Wilcoxon Rank Sum test **** p<<0.0001). **C** Ongoing activity in unmatched cells in DDD mice at ZT0 and ZT2 (cells recorded from 4 mice; ZT0: n = 2564 cells; ZT2 dark: n = 2364 cells; Statistics Wilcoxon Rank-Sum test **** p<<0.0001). **D** Representative traces of daily changes in ongoing activity in a representative longitudinally tracked neuron (green traces (F) = dF/F signals; black traces (D) = corresponding deconvolved signals). **E** Daily changes in ongoing activity in all matched cells (n = 1148 cells, recorded from 7 mice; (ZT0 vs ZT2 p = 1.22e-25, ZT0 vs ZT12 p = 0.23, Wilcoxon Signed-Rank test) **F,G** Daily changes in ongoing activity of ‘Ever’ (**F**) or ‘Stable’ (**G**) visually responsive cells (cells recorded from 4 mice; ‘Ever’ cells: n= 840 cells; ‘Stable’ cells: n= 244 cells; Statistics Wilcoxon Signed-Rank **** p<0.0001). **H-J** Direction tuning, orientation selectivity and preference of individual neurons are stable across the day. Direction tuning curves of ‘Ever’ (**H**) or ‘Stable’ (**I**) visually responsive cells across the day (ZT0 = red, ZT2 = green, ZT12 = blue; diamonds = normalized average responses; lines = normalized double-gaussian-fitted population averages). Insets: Global orientation selectivity indices (gOSI) of respective cells at ZT0, ZT2, or ZT12 show no significant differences across the day (n.s. = not significant). **J** Orientation preference of ‘Stable’ cells does not change across the day. Left: fitted preferred orientation of individual neurons between ZT0 and ZT2 (top) or ZT12 (bottom). Right: Histograms showing difference in fitted preferred orientation between ZT0 and ZT2 (top) or ZT12 (bottom). (Error bars represent SEM in all data panels; see **Supplemental Data Table 3** for descriptive statistics)

We first analyzed the overall ongoing activity of all imaged neurons, i.e. the activity of all neurons in the field of view (FOV) when no visual stimulus is presented to the mice and without comparing the activity of single neurons across the imaging sessions (i.e. the cells were not “matched” between sessions). Strikingly, we found that the activity of all the neurons in the FOV (i.e. population activity) increases two hours after light onset at ZT 2 (increase of 22.5%) and that this increase is largely gone ten hours later at ZT 12 (**Figure 3B**). Importantly, we did not observe such an increase in the ongoing activity of visual cortex L2/3 excitatory neurons in DDD mice (**Figure 3C**), thus demonstrating that it is the daily appearance of light that causes the initial rapid increase and subsequent normalization in the ongoing activity of the imaged neurons.

Having identified a daily light-driven increase and normalization in ongoing activity in L2/3 PYR neurons at the population level, we next asked whether this occurs also in “matched” single neurons, i.e. in L2/3 PYR neurons that we were able to reliably identify and track their activity in all three imaging sessions across the day and to compare the daily changes in the activity of each neuron to itself (“Matched cells”; **Figure 3D**). This revealed that the activity of individual neurons increases at ZT 2 by 31.5% on average (**Figure 3E**). Notably, these daily light-driven changes in ongoing activity in “matched” neurons were not caused by different behavioral states (e.g. arousal) of the animals at the different times of the day: using pupil-dilation (**Supplemental Figure 3A**) and whisking (**Supplemental Figure 3B**) as two independent and reliable indicators of the animals’ arousal states ^63,64^, we found that these daily light-driven changes in ongoing activity occur regardless of whether the animals are in low or high arousal states.

Finally, we asked whether the daily appearance of light alters also the visual response properties of L2/3 excitatory neurons in the visual cortex. For this, we focused on matched neurons that responded to visual stimuli during at least one of the imaging sessions (i.e. at ZT 0, 2 and/or 12) and analyzed two groups of neurons: those that responded to visual stimulation at least at one time point (“Ever” cells, 43.2% of all matched cells) and those that responded at all three timepoints (“Stable” cells, 29.6% of all “Ever” cells). As expected - and similar to our observations in “unmatched” and “matched” neurons (**Figure 3B,E**) - we found that ongoing activity transiently increases at ZT 2 both in “Ever” and in “Stable” neurons (**Figure 3F,G**). However, when we analyzed visual response properties such as direction tuning, orientation selectivity, spatial frequency tuning and contrast sensitivity at the three timepoints, we did not observe any changes in these properties across the day in either group of cells (**Figure 3H-J** and **Supplemental Figure 3C-H**).

Taken together, these experiments reveal that the daily appearance of light causes every day a rapid and transient increase in the ongoing activity of visual cortex excitatory neurons without changing their tuning properties. These findings are consistent with our observation that E/I-ratio in these neurons transiently shifts towards more excitation one hour after the appearance of light and that E/I-ratio in these neurons is normalized several hours later by the light-driven increase in inhibitory inputs (**Figure 2**). Since many of the daily light-induced genes identified in our transcriptomic analyses (**Figure 1**) were previously found to increase inhibition onto principal excitatory neurons in the neocortex in a cell-autonomous manner (e.g *Npas* and *Fos* onto CA1 pyramidal neurons ^50,51,65^, *Bdnf* onto cortical neurons ^45,52,66,67^) and since activity-induced transcriptional networks exert their biological functions several hours after stimulus onset ^28,33,34^, this raises the possibility that daily light-induced transcriptional programs drive every day the light-dependent increase in inhibitory inputs onto visual cortex excitatory neurons to normalize E/I-ratio and neural firing rates in these neurons and to thereby maintain the tuning properties of these neurons over time.

### NPAS4 promotes every day inhibitory inputs onto visual cortex neurons in a light-dependent manner to maintain E/I-ratio

NPAS4 is a ubiquitous early-induced transcription factor that promotes inhibitory inputs onto neocortical excitatory neurons in a cell-autonomous manner by activating cell-type-specific sets of late-induced effector genes ^38,50,51,68^. Since the daily appearance of light induces *Npas4* in visual cortex excitatory neurons across all cortical layers (**Figure 1D** and **Supplemental Figure 1G**) we hypothesized (1) that NPAS4 is part of an activity-induced genetic network that normalizes the rapid daily light-induced shift in E/I-ratio towards excitation and concomitant increase in neural activity in visual cortex excitatory neurons by driving the slower daily light-induced increase in inhibitory inputs, and (2) that NPAS4 thereby maintains proper sensory processing in the visual cortex.

To test this idea, we first established a loss-of-function approach to knock out *Npas4* selectively in excitatory neurons in the adult visual cortex. For this, we generated two novel AAV constructs that drive the expression of either an active or an inactive form of Cre-recombinase (Cre and dCre, respectively) and of the red-fluorescent protein mRuby3 in cortical excitatory neurons (**Supplemental Figure 4A**), injected these AAVs into the visual cortices of mice that are homozygous for an allele of *Npas4* that is flanked by LoxP sites and that can be removed by Cre-expression (i.e. floxed *Npas4* allele ^50^), housed these mice in the dark for two weeks, exposed them to light for two hours and then immunolabeled visual cortex sections derived from these mice with antibodies directed against NPAS4. These experiments revealed that NPAS4 is absent (i.e. “knocked out”, cKO) in neurons transduced with the Cre-expressing AAV construct (**Supplemental Figure 4B-D**).

We then used this AAV-based knockout approach to test whether NPAS4 promotes inhibitory inputs onto L2/3 excitatory neurons in the adult visual cortex in an acute light-dependent manner. Since we wanted to test whether NPAS4 exerts its function in these neurons in a cell-autonomous manner, we injected the respective AAV construct (Cre for *Npas4* cKO or dCre as control) at considerably lower concentrations than in our calibration experiments such that the AAV-infected neurons (less than 10% of all excitatory neurons, see Methods) are surrounded by non-infected (i.e. *Npas4* wild-type) neurons and that the observed effects are due to the cell-autonomous effects of *Npas4* knockout in the infected neurons but not due to potential network effects. To identify the short-term (i.e. acute) sensory-dependent cellular functions of NPAS4, we housed the mice after AAV injection in the dark for seven to ten days, then prepared acute visual cortex slices either without exposing the mice to light or after twelve hours of light-exposure, and used whole cell patch-clamp electrophysiology to record mIPSCs and mEPSCs in AAV-infected L2/3 PYR neurons and to determine E/I-ratio in these neurons (**Figure 4A**). This revealed that in control-infected L2/3 PYR neurons, light-exposure after dark-housing led to an increase in inhibitory inputs (i.e. an increase in mIPSC frequency and amplitude upon light-exposure) but not to a light-induced change in E/I-ratio since also the excitatory inputs onto these neurons increased in a light-dependent manner (i.e. mEPSC frequency increases) (**Figure 4B,C**). However, upon cKO of *Npas4*, the appearance of light caused a shift in E/I-ratio towards excitation (**Figure 4C**) since the light-induced increase in inhibitory inputs did not occur while the light-induced increase in mEPSC frequency still occurred (**Figure 4B**). Since (1) *Npas4* is largely absent in visual cortex excitatory neurons in dark-housed mice and is expressed only upon light-exposure (**Supplemental Figure 4E**), and (2) we did not observe significant changes in mEPSCs, mIPSCs and E/I-ratio in the dark-housed mice, we conclude that the acute/short-term cellular function of NPAS4 in visual cortex L2/3 PYR neurons lies in the light-induced promotion of inhibitory inputs onto these neurons and the maintenance of E/I-ratio.

**Figure 4.**
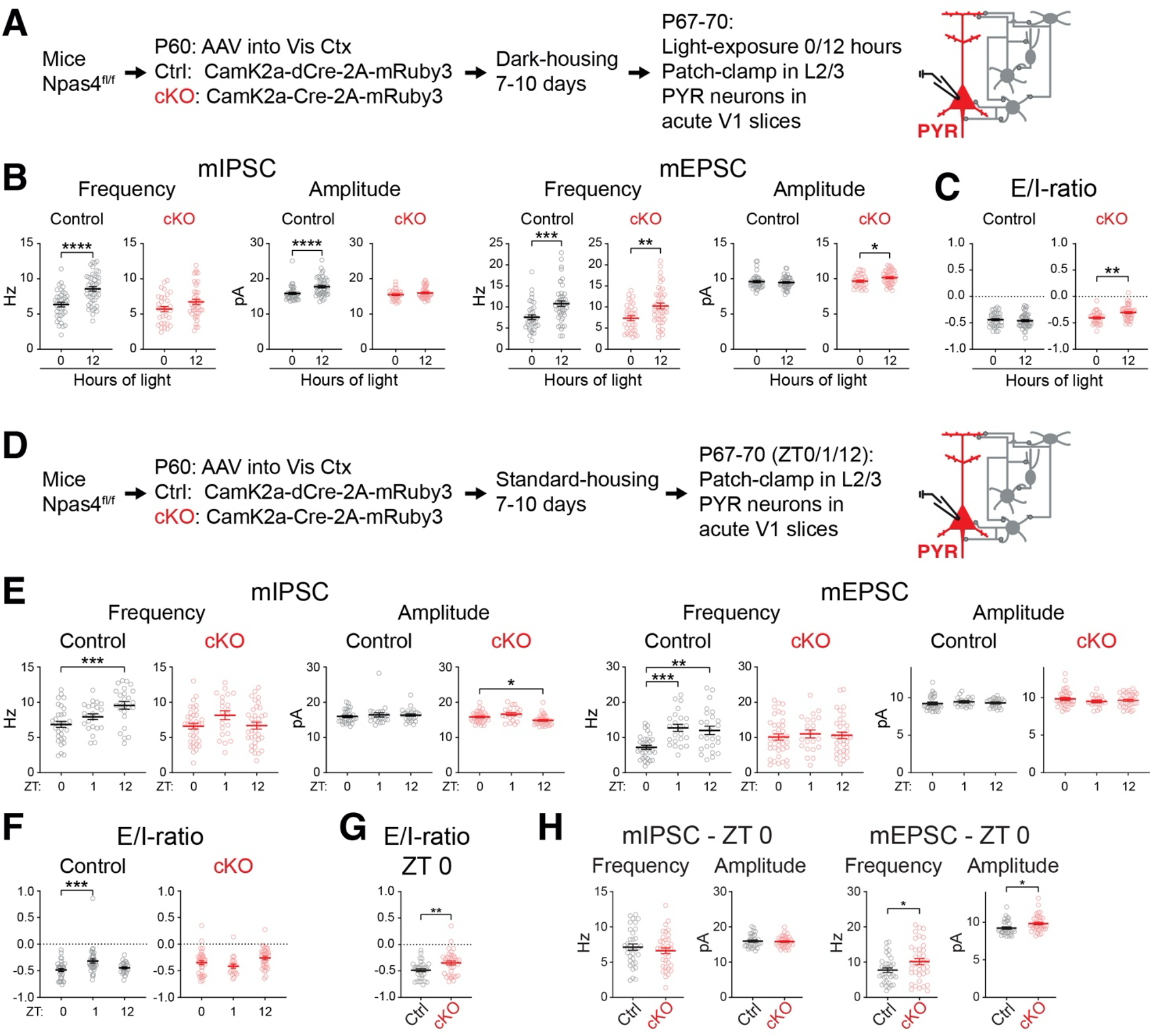
NPAS4 promotes every day inhibitory inputs onto visual cortex neurons in a light-dependent manner to maintain E/I-ratio. **A-C** NPAS4 controls sensory-dependent plasticity of inhibitory inputs onto visual cortex L2/3 PYR neurons and thereby maintains E/I-ratio. **A** Experimental strategy for whole-cell patch-clamp recordings from L2/3 PYR neurons in acute visual cortex slices in the context of a dark-housing/light-exposure sensory stimulation paradigm and upon AAV-mediated knockout of *Npas4* in excitatory neurons in the adult visual cortex (AAV-infected cells are identified by mRuby3 expression; cKO - Cre-2A-mRuby3, Control - dCre-2A-mRuby3). **B** Quantified data of frequencies and amplitudes of mIPSCs, mEPSCs and E/I-ratio of *Npas4* control (black) and cKO (red) neurons (Control: 0 hours of light n = 37 cells from 4 mice, 12 hours of light n = 46 cells from 4 mice; cKO: 0 hours of light n = 31 cells from 4 mice, 12 hours of light n = 42 cells from 5 mice; Statistics Mann-Whitney * p<0.05, ** p<0.01, *** p<0.001, **** p<0.0001). **C** Quantified data of E/I-ratio of *Npas4* control and cKO neurons (Statistics Mann-Whitney ** p<0.01). **D-H** NPAS4 drives that daily light-induced increase in inhibitory inputs and in the absence of Npas4, E/I-ratio shifts over time towards excitation. **D** Experimental strategy for whole-cell patch-clamp recordings from L2/3 PYR neurons in acute visual cortex slices in the context of standard-housing and upon AAV-mediated knockout of *Npas4* in excitatory neurons in the adult visual cortex (AAV-infected cells are identified by mRuby3 expression; cKO - Cre-2A-mRuby3, Control - dCre-2A-mRuby3) **E, F** Quantified data of frequencies and amplitudes of mIPSCs and mEPSCs (**E**) and E/I-ratio (**F**) of *Npas4* control (black) and cKO (red) neurons (Control: ZT0 n = 34 cells from 4 mice, ZT1 n = 37 cells from 4 mice, ZT12 n = 25 cells from 4 mice; cKO: ZT0 n = 37 cells from 4 mice, ZT 1 n = 21 cells from 4 mice, ZT12 n = 34 from 4 mice; Statistics Kruskal-Wallis test with post-hoc Dunnett’s T3 multiple comparisons test, * p<0.05, ** p<0.01, *** p<0.001, **** p<0.0001). **G, H** Direct comparison of E/I-ratio (**G**) and frequencies and amplitudes of mIPSCs and mEPSCs (**H**) in control and cKO neurons at ZT0 (Control: n = 34 cells from 4 mice; cKO: ZT0 n = 37 cells from 4 mice; Statistics Kruskal-Wallis test with post-hoc Dunnett’s T3 multiple comparisons test, * p<0.05, ** p<0.01). (Error bars represent SEM in all data panels; see **Supplemental Data Table 2** for descriptive statistics)

Next, we asked about the longer-term functions of NPAS4 in visual cortex L2/3 PYR neurons, particularly whether NPAS4 drives the daily normalization of E/I-ratio in these neurons via the daily light-induced increase in inhibitory inputs. For this, we took the same sparse loss-of-function approach as in our short-term stimulation experiments (**Figure 4A,B**) but instead of dark-housing the AAV-injected mice, we housed the mice in a regular 12/12 dark-light cycle (**Figure 4D**) and measured mIPSCs, mEPSCs and E/I-ratio at ZT 0, ZT 1 and ZT 12. This revealed that the daily light-driven increase in excitatory and inhibitory inputs (**Figure 4E**) and normalization of E/I-ratio (**Figure 4F**) occurred in control-infected L2/3 PYR neurons similar as in wild-type mice (**Figure 2**) but that upon *Npas4* knockout, inhibitory inputs and E/I ratio were not modulated across the day and excitatory inputs underwent only weak strengthening. Since NPAS4 promotes inhibitory inputs onto L2/3 PYR neurons in an acute light-dependent manner to maintain E/I-ratio (**Figures 4A-C**), we next compared the E/I-ratio in knockout and control neurons at ZT 0: this revealed that the E/I-ratio in *Npas4* knockout neurons is indeed shifted towards excitation (**Figure 4G**). Interestingly, this shift in E/I-ratio towards excitation upon *Npas4* knockout is caused by an increase in excitatory inputs (**Figure 4H**) which likely is a secondary, longer-term effect of lacking NPAS4 in these neurons when mice are housed under standard dark-light conditions where *Npas4* is normally induced every day to normalize E/I-ratio. Thus, taken together, our experiments reveal (1) that the light-induced expression of *Npas4* in visual cortex L2/3 PYR neurons promotes inhibitory inputs onto these neurons within several hours after the appearance of light, and (2) that NPAS4 expression in these neurons maintains E/I-ratio by preventing a shift towards more excitation.

### NPAS4 normalizes activity rates in visual cortex excitatory neurons every day and maintains orientation tuning over time

Having established the cellular-synaptic effects of *Npas4* knockout in visual cortex L2/3 PYR neurons, we then set out to test whether these effects - particularly the lack of light-induced normalization of E/I ratio (**Figure 4C**) - affect the daily normalization of the light-induced increases in neural activity in these neurons and their visual response properties. For this, we took a similar genetic approach as in our electrophysiology cKO experiments: we co-injected d/Cre-2A-mRuby3 AAVs together with an AAV that expresses GCaMP7f in excitatory neurons into mV1 of adult *Npas4* flox mice, housed the AAV-injected mice for two to three weeks in a standard 12/12 hours dark-light cycle (we acclimated the mice to the imaging setup and head-fixation during this period, see Methods) and analyzed the neural activity and visual response properties in AAV-transduced L2/3 PYR neurons via *in vivo* 2-photon microscopy at ZT0, ZT2 and ZT12 (**Figure 5A,B**). We restricted our analyses in these experiments to “matched” cells that co-expressed GCaMP7f and mRuby3, as this would allow us to identify in single cells daily changes in ongoing activity and/or response properties that were caused by the cKO of *Npas4* in these cells.

**Figure 5.**
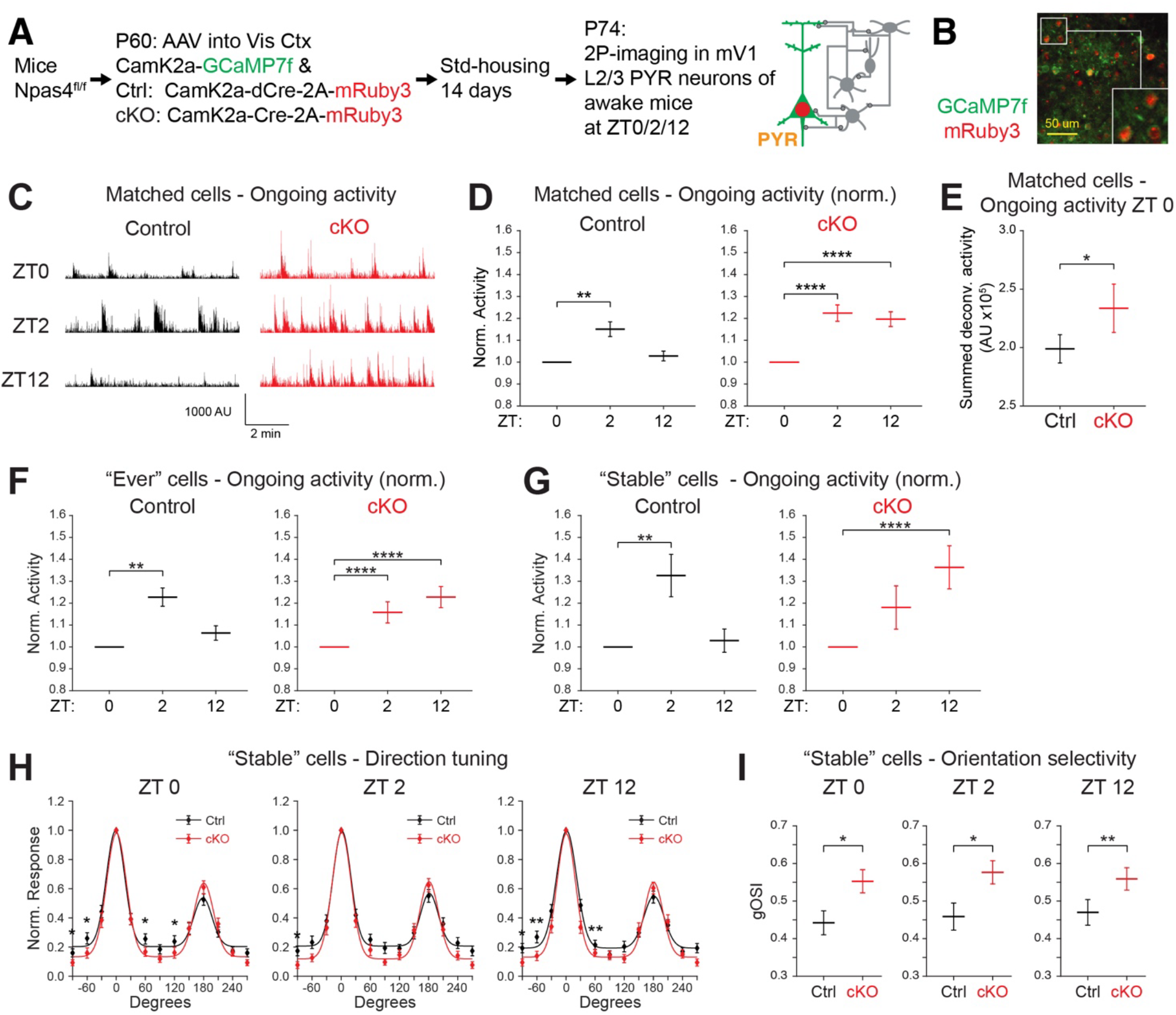
NPAS4 normalizes activity rates every day in visual cortex excitatory neurons every day and maintains orientation tuning over time. **A** Experimental strategy to analyze ongoing activity and response properties in visual cortex L2/3 excitatory neurons in standard housed mice upon AAV-mediated knockout of *Npas4* (AAV-infected cells are identified by mRuby3 expression; cKO - Cre-2A-mRuby3, Control - dCre-2A-mRuby3). **B** Representative image of excitatory neurons in mV1 L2/3 of *Npas4* fl/fl mice that express GCaMP7f (green) and mRuby3 after stereotactic injection of AAVs (Scale-bar = 50 micron). The subsequent analyses of neural activity were based on cells co-expressing GCaMP7f and mRuby3. **C-G** NPAS4 normalizes the daily light-induced increase of ongoing activity in excitatory neurons every day. **C** Representative traces of ongoing activity (deconvolved signal) from one longitudinally tracked *Npas4* cKO (red) and Control (black) neuron at ZT0, ZT2, and ZT12. **D** Daily changes in ongoing activity in matched *Npas4* cKO (red) and Control (black) neurons (n = Ctrl: 400 neurons from 4 mice, cKO: 500 neurons from 5 mice [100 random neurons/mouse]; Wilcoxon Signed-Rank test ** p<0.01). **E** Ongoing activity at ZT0 in *Npas4* Control (black) and cKO (red) neurons (n = Ctrl: 400 neurons from 4 mice, cKO: 500 neurons from 5 mice [100 random neurons/mouse]; Wilcoxon Rank Sum test * p<0.05). **F, G** Daily changes in ongoing activity in ‘Ever’ (**F**) or ‘Stable’ (**G**) visually responsive neurons that are cKO (red) or Control (black) for *Npas4* (‘Ever’ neurons: cKO - n = 365 cells from 4 mice, Ctrl - n = 388 cells from 4 mice; ‘Stable’ neurons: cKO - n = 108 cells from 4 mice, Ctrl - n = 71 cells from 4 mice; Wilcoxon Signed-Rank test * p<0.05, ** p<0.01, **** p<0.0001). **H, I** NPAS4 maintains orientation tuning in visual cortex excitatory neurons. Direction tuning curves (**H**) and global Orientation Selectivity Index (gOSI) (**I**) of ‘Stable’ visually responsive neurons that are cKO (red) or Control (black) for *Npas4* at ZT0, ZT2 and ZT12 (Control n = 42 cells, cKO n = 40 cells; Wilcoxon Rank Sum test * p<0.05, ** p<0.01). (Error bars represent SEM in all data panels; see **Supplemental Data Table 3** for descriptive statistics)

Using this experimental setup, we first analyzed whether the daily light-induced increase and subsequent normalization of ongoing activity still occurs in *Npas4* knockout and control neurons (**Figure 5C-E**). This revealed that the daily rapid light-driven increase in ongoing activity at ZT 2 occurs in control and *Npas4* knockout neurons similar as in wild-type neurons (**Figure 3**) but that the subsequent normalization of ongoing activity occurs only in control-infected neurons while ongoing activity remains elevated in *Npas4* cKO neurons (**Figure 5C,D**). Consistent with this finding and with our observation that the lack of *Npas4* causes a shift in E/I-ratio toward excitation already at ZT 0 (**Figure 4C,G**), we also found that *Npas4* cKO neurons are more active already at ZT 0, indicating that the baseline activity of these neurons increases when *Npas4* is absent for multiple days (**Figure 5E**). Notably, this lack of daily normalization of ongoing activity upon *Npas4* knockout is not specific to “Stable” or “Ever” cells, and also the proportion of these populations among all matched cells is only marginally affected by *Npas4* cKO (Control: 31.3% “Stable” cells; cKO: 30.0% “Stable” cells).

Having established that NPAS4 is required for the daily normalization of ongoing activity, we then asked whether the lack of *Npas4* in visual cortex L2/3 PYR neurons affects their visual response properties. For this, we analyzed “Stable” and “Ever” cells separately and first focused on direction tuning and orientation selectivity. These analyses revealed that the lack of *Npas4* causes in “Stable” and “Ever” cells a similar increase in orientation selectivity (**Figure 5G,I** and **Supplemental Figure 5A,B**) and that these cKO-induced changes are not subject to daily modulation (i.e. the orientation selectivity of each cKO population is altered to a similar degree at ZT 0, 2 and 12). Interestingly, the effects of *Npas4* cKO on the tuning properties of L2/3 PYR neurons are selective as we did not observe any changes in the contrast sensitivity of either neuronal population upon *Npas4* cKO (**Supplemental Figure 5C,D**), and a partial impairment in spatial frequency tuning only in “Stable” cells (**Supplemental Figure 5E,F**). Thus, taken together, these experiments reveal that the daily light-induced expression of *Npas4* in visual cortex excitatory neurons is necessary for the daily normalization of the daily light-induced increase in ongoing neural activity in these neurons and for stabilizing their orientation tuning. Since the activity and tuning properties of neurons affect an animal’s sensory perception ^69^, these findings suggest that the daily light-induced expression of *Npas4* in visual cortex excitatory neurons maintains stable visual perception over time.

## Discussion

Homeostatic control over neural activity rates is thought to be essential for maintaining an animal’s ability to properly process, store and perceive (sensory) information ^11^ but it remains unknown whether such Firing Rate Homeostasis (FRH) occurs during an animal’s everyday life, and the molecular mechanisms that exert FRH *in vivo* in intact neural circuits are poorly understood. Since FRH occurs in response to experience-driven changes in neural activity and counteracts other experience-induced changes in synaptic weights (e.g. Hebbian strengthening of synapses), the molecular mechanisms that exert FRH must themselves be activity-regulated and exert their cellular and circuit functions in time-spans that match those of FRH (i.e. over several hours). However, while multiple genes and molecules have been implicated in regulating homeostatic forms of cellular-synaptic plasticity e.g. in cultured neurons ^70^, only a subset of these molecular mechanisms were demonstrated to be experience-regulated *in vivo* in intact neural circuits on timescales relevant for controlling FRH, and only one these genes and molecules was shown to control FRH in single neurons *in vivo* in the brain (the synaptic scaffolding molecule SHANK3 is necessary for upward FRH in the developing visual cortex in response to MD ^32^). Here, we combined in one coherent experimental setting a quasi-naturalistic sensory stimulus with transcriptomic, cellular-synaptic and circuit-level analyses and with a targeted genetic manipulation in the adult cortex to demonstrate - to our knowledge for the first time - that (1) the daily appearance of light leads in PYR neurons in the adult visual cortex to a rapid increase in excitatory inputs and shift in E/I-ratio towards excitation and to a rapid concomitant increase in the activity rates of these neurons, and (2) that light-induced transcription via the early-induced TF NPAS4 subsequently promotes inhibitory inputs onto these neurons to return the E/I-ratio and neural activity rates of these neurons to their pre-stimulus levels (i.e. to their levels before the appearance of light) and to maintain proper orientation tuning in these neurons (**Figure 6**). Thus, our study reveals that daily light-induced transcription in PYR neurons in the adult visual cortex normalizes E/I-ratio to drive downward FRH and to maintain proper sensory processing. These findings are noteworthy in several regards.

**Figure 6:**
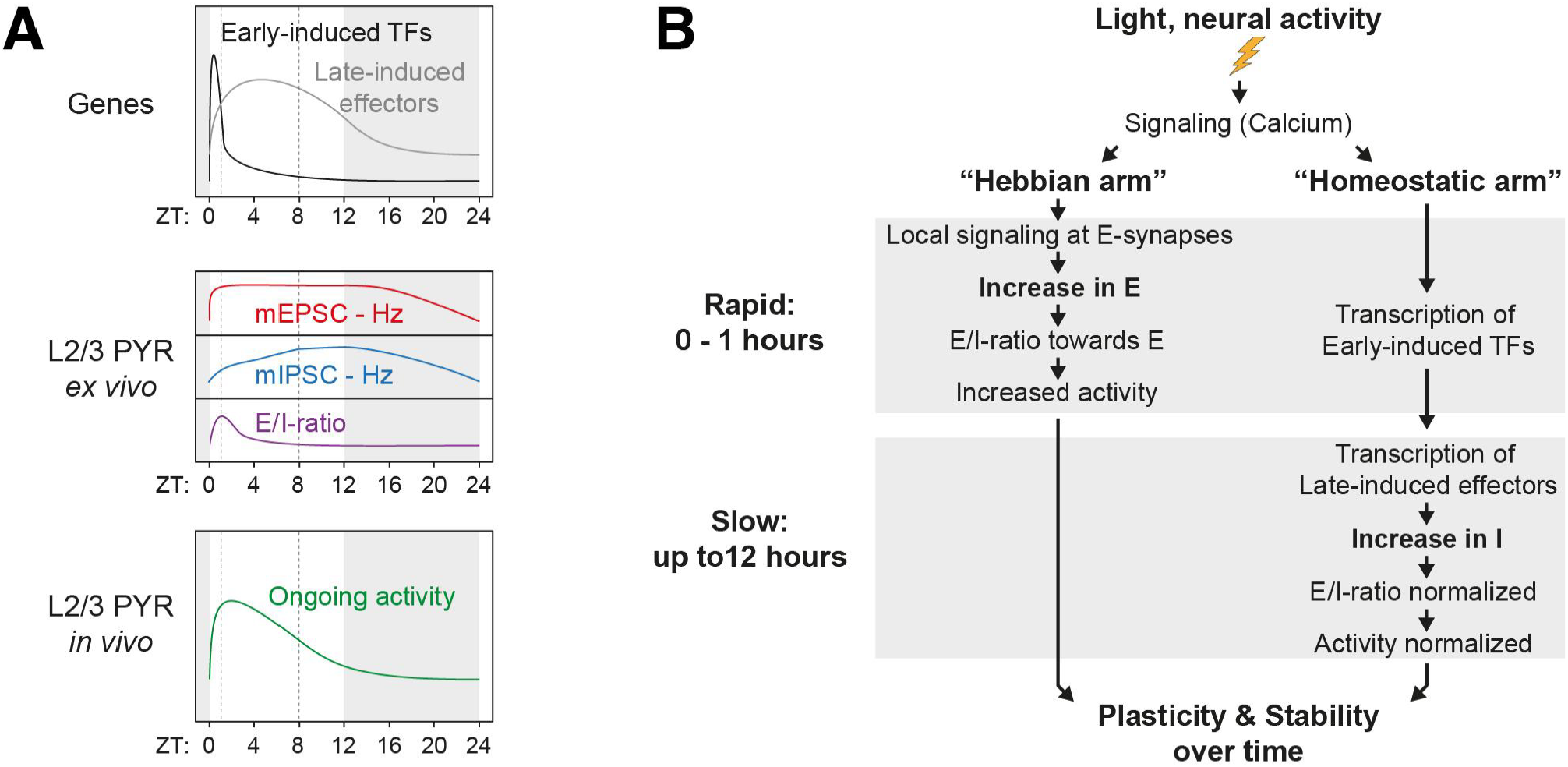
Light-induced transcription maintains firing rate homeostasis and visual function by normalizing E/I-ratio every day. **A** Summary of daily light-induced effects on transcription, excitatory and inhibitory inputs, E/I-ratio and ongoing activity in visual cortex neurons. **B** A model for how one sensory stimulus (e.g. the daily appearance of light) can induce two temporally coordinated yet opposing molecular and cellular cascades in the same neuron (e.g. in visual cortex excitatory neurons): the more rapid process drives within one hour after stimulus onset a rapid increase of excitatory inputs (e.g. via transcription-independent events) and shifts E/I-ratio towards excitation while the slower process drives an increase of inhibitory inputs over several hours, thereby normalizing E/I-ratio and neural activity rates in a homeostatic manner and maintaining neural circuit function over time. Together, these two activity-induced forms of synaptic modulation convey both plasticity and functional stability to the neural circuits such as the adult visual cortex where activity-induced transcriptional networks are essential for stabilizing FRH and sensory perception every day.

Our experiments clearly demonstrate that it is the daily appearance of light (i.e. a sensory experience) *per se* that drives the daily transcriptomic, cellular-synaptic and circuit changes in visual cortex neurons and that these daily light-induced changes occur in the absence of specific plasticity-/learning paradigms and independent of events that may co-occur with the appearance of light on the same day (e.g. changes driven by circadian mechanisms) or light-induced changes in the animals’ behavioral states (**Supplemental Figure 3A,B**). Notably, the time courses of these light-induced changes in gene transcription, synaptic modulations and changes in neural activity are highly consistent with each other (see **Figure 6A**), thus further strengthening the validity of our findings. Also, our study reveals that the large neuron-to-neuron variability in the activity rates of visual cortex neurons which was observed by us (**Figure 3**) and others ^71^ necessitates analyzing a large number of neurons (i.e. more than one hundred) to identify these daily light-induced changes in the activity of visual cortex (PYR) neurons (**Supplemental Figure 3I**). This might explain why previous studies based on longitudinal electrophysiological recordings in the rodent visual cortex did not observe similar daily light-induced changes in the activity of visual cortex neurons ^17,72,73^: in addition to the animals’ different ages in these studies (3-5 weeks old versus 8-10 weeks old in our experiments) and different stimulation paradigms used in some of them ^17^, these studies analyzed only relatively small numbers of neurons (i.e. several dozen neurons) and were thus underpowered to detect the daily light-induced changes in neural activity that we identified in our current study. Our study further reveals that detecting these daily light-induced increases in neural activity requires that the recordings are done relatively shortly after the appearance of light - this might explain why previous studies based on calcium imaging of visual cortex neurons did not observe daily changes in neural activity: while the number of neurons analyzed in these studies were sufficient to detect such daily changes, the timing of the recordings in these studies relative to the stimulus onset was rather different from the schedule that we used in our current study and was therefore not suitable for detecting daily light-induced changes in neural activity ^14,16^. Thus, these technical differences are likely to explain why previous studies in the rodent visual cortex did not observe this naturally occurring form of downward FRH that we report on here.

Consistent with our *in vivo* calcium imaging experiments, our *ex vivo* electrophysiological recordings in acute visual cortex slices clearly establish that the daily appearance of light modulates excitatory and inhibitory inputs onto visual cortex L2/3 PYR neurons on distinct time scales. The net result of these synaptic modulations is that the appearance of light rapidly shifts E/I-ratio towards excitation which is followed by a slower shift in E/I-ratio back towards inhibition such that E/I-ratio is normalized towards the end of the day’s light phase. Our use of established approaches ^60,74^ to measure mEPSCs and mIPSCs in the same cells by switching the holding potential of the cells without using synaptic blockers was essential to precisely determine the E/I-ratio in each cell and to detect the light-induced changes in E/I-ratio across the population of recorded neurons. Thus, our technical approach differed from the one used in a previous study ^75^ where mEPSCs and mIPSCs were isolated pharmacologically and E/I-ratio could be calculated only based on mEPSCs and mIPSCs that were recorded in separate neurons. This technical difference matters since the frequency of mEPSCs and mIPSCs strongly varies between individual cells (**Figure 2**) and the calculation of population average E/I-ratio based on population averages of mEPSCs and mIPSCs may yield different values than when calculated based on the E/I-ratios of individual cells as done in our study.

Our findings on the cellular-synaptic functions of the early-induced TF NPAS4 are consistent with previous observations by us ^38^ and others ^50,51,76,77^: similar as e.g. in PYR neurons in the CA1 region of the mouse hippocampus ^50,51,76,77^, we find that the acute/short-term cellular function of NPAS4 in visual cortex PYR neurons lies in the cell-autonomous sensory-induced promotion of GABAergic inputs and homeostatic maintenance of E/I-ratio. Our experiments further reveal that the longer-term function of NPAS4 in visual cortex PYR neurons lies in maintaining E/I-ratio as well but that the underlying cellular mechanisms seem to change over time (**Figure 4**). However, our experiment cannot determine whether the increase in excitatory inputs upon longer-term knockout of *Npas4* is due e.g. to cell-autonomous mechanisms based on altered expression of genes that are not directly regulated by NPAS4 and/or due to circuit effects that come to bear over time despite the relatively sparse knockout of *Npas4* in visual cortex excitatory neurons - future experiments will have to test these possibilities. Similarly, future experiments will have to determine the type(s) of GABAergic inputs onto L2/3 PYR neurons that are regulated by NPAS4 in visual cortex PYR neurons: given that the daily appearance of light induces the transcription of multiple early-induced TFs in these neurons and that experiments in hippocampal CA1 PYR neurons revealed that NPAS4 and FOS promote different sets of somatic GABAergic inputs ^65,76^, it is tempting to speculate that TF-specific transcriptional subnetworks modulate every day different sets of inhibitory inputs onto visual cortex PYR neurons. In turn, this raises the exciting possibility that functional analyses of these activity-induced transcriptional subnetworks will allow for manipulating sensory-induced plasticity in specific (GABAergic) inputs onto PYR neurons to thereby determine the daily impact of these input-specific plasticity mechanisms on the activity and response properties of these neurons. An alternative, not mutually exclusive explanation for such light-induced expression of multiple early-induced TFs in the same neurons is that these genes can compensate for each other to thereby prevent potentially catastrophic consequences of even subtle genetic mutations in these genetic networks. Future experiments will have to test these ideas.

Our finding that NPAS4 drives downward FRH in an experience-dependent manner is consistent with the role of this early-induced TF in the homeostatic maintenance of E/I ratio (**Figure 4** and ^38,50,51,76,77^). Thus, while the finding that NPAS4 drives downward FRH might not be surprising, it is important nevertheless since it provides - to the best of our knowledge - the first identification of an activity-induced molecular mechanism that drives downward FRH *in vivo*. Our observation that NPAS4 is necessary for maintaining select response properties in visual cortex PYR neurons is in this regard of particular interest: as was suggested by theoretical studies ^8,11^, this finding suggests that the maintenance of FRH and of proper response properties are mechanistically connected and future experiments will have to test this idea in depth. Similarly, while we did not assess whether the changes in orientation tuning caused by *Npas4* knockout in visual cortex PYR neurons alters an animal’s visual perception, one might expect that this is the case: similar changes in the orientation tuning of visual cortex PYR neurons were previously found to affect an animal’s ability to discriminate between differently oriented gratings ^78^, and future experiments thus will have to test whether the daily light-induced expression of NPAS4 in visual cortex PYR neurons is necessary to maintain an animal’s visual perception. Furthermore, since recent experiments in hippocampus PYR neurons indicate that activity-induced transcription via the early-induced TF FOS restricts representational drift in neural circuits ^79^ and the stability of engrams ^80^, an interesting idea is that NPAS4 exerts similar functions. Thus, our study provides a conceptual and experimental framework for future studies in which *Npas4* knockout in adult cortical circuits can be used to explore the interrelations between FRH, sensory processing, perception, representational stability and the formation and maintenance of engrams.

Lastly, an important question is whether similar homeostatic mechanisms based on experience-induced transcription maintain the functional stability of neural circuits other than the visual cortex. Given that experience-induced gene transcription was observed in many neural circuits ^33,34,36,81–85^ and that experiments e.g in the hippocampus, the prefrontal cortex and the somatosensory cortex revealed that early-induced TFs such as NPAS4 ^48,86,87^ and FOS ^65^ and late-induced effectors such as ARC ^88^, NARP ^53^ and BDNF ^51^ regulate specific sets of synapses and E/I-ratio according to a homeostatic circuit logic also in these neural circuits, it seems likely that this is indeed the case. We therefore propose that a generalizable function of activity-induced transcription in neural circuits lies in the activity-induced normalization of E/I-ratio and control over FRH to thereby counterbalance other forms of experience-dependent cellular-synaptic plasticity (e.g. learning-associated Hebbian plasticity) and to maintain proper response properties and stabilize perception over time.

## Acknowledgments

We thank all the members of the Spiegel lab for comments and discussions, and Mr. Hasan Heidar and Ms. Shakked Ganor for assistance with animal husbandry. This work was supported by an ISF I-CORE grant (1916/12), an ISF personal grant (2354/19) a BSF US-Israel binational grant (2017342), all for IS. DK is supported by a fellowship from the Israel Ministry of Absorption (IMOAb) and by the Horowitz Foundation. DA is supported by a fellowship from the Ariane de Rothschild Women Doctoral Program. IS is the incumbent of the Friends and Linda and Richard Price Career Development Chair and a scholar in the Zuckerman STEM leadership program.

## Author contribution

ET, DK and IS initiated and conceived this project; DK conducted and analyzed all the experiments in Figures 3 and 5 and in Supplemental Figures 3 and 5; ET conducted all the experiments in Figures 2 and 4 and in Supplemental Figures 2 and 4; DA conducted and analyzed the experiments in Figure 1A-C and Supplemental Figure 1A-F; OR conducted the experiment in Supplemental Figure 1H and helped ET with cloning of the AAV constructs described in Supplemental Figure 4A and MB-K with the experiments in Figure 1D,E and Supplemental Figure 1G; MBK conducted and analyzed the experiments in Figure 1D,E and Supplemental Figure 1G; KCK oversaw and contributed to the analyses of the experiments in Figures 2-5 and Supplemental Figures 2-5; IS generated Figure 6, acquired funding, supervised the project and wrote the manuscript with input from all authors.

## MATERIALS & METHODS

### Animals and Dark-Housing/Light-Exposure (DH/LE) sensory stimulation paradigm

All procedures involving animals were reviewed and approved by the Weizmann Institutional Animals Care Committee. Experiments were done in young adult mice of either sex, housed up to five animals per cage in a 12/12 dark/light cycle. All mice used in this study were WT or homozygous for any cKO allele. Animals were housed in standard circadian cabinets to precisely time light on- and offset.

For DH/LE sensory stimulation, mice were dark housed for a minimum of 7 days in light sealed cabinets and the cages were refreshed once after 3-5 days in the dark using night goggles. Light-exposure was performed in the same cabinets with bright white LEDs situated above the cage for one or three hours after which the mice were sacrificed and the tissue was dissected. Animals that were not light-stimulated were enucleated in the dark with night goggles prior to dissecting the tissue.

### Cloning of DNA constructs

AAV plasmid constructs were generated using standard cloning techniques, and the integrity of all cloned constructs was validated by DNA sequencing. To generate the pAAV-CW3SL-mRuby3-2A-Cre-WPRE and pAAV-CW3SL-mRuby3-2A-dCre-WPRE constructs, we used pAAV-CW3SL-EGFP (Addgene plasmid #61463) as a backbone and replaced EGFP with PCR products that contained either mRuby3-2A-Cre-WPRE or mRuby3-2A-dCre-WPRE (the plasmids pAAV-Ef1a-fDIO-mRuby3-2A-Cre and pAAV-Ef1a-fDIO-mRuby3-2A-dCre respectively served as templates ^28^) via restriction with EcoRI and NheI.

### AAV production

AAV particles were produced as previously described ^28,89^. Briefly, HEK293T (ATCC) cells were grown in DMEM supplemented with 5% FBS, penicillin-streptomycin (Life technologies), NEAA (11140050, Gibco™) and sodium pyruvate (1mM; 03-042-1B, BI) and were plated the day before transfection at a density of 12 million cells on 15cm poly-L-lysine coated plates. One day later, equal amounts of plasmids containing sequences for pDJ, pHelper, and the desired expression cassette (13.3μg per plasmid) were transfected onto plates using PEI. Two days post transfection, the medium was collected and stored at 4°C and replaced with fresh medium. Five days post transfection, the cells and medium were collected for viral purification. AAV particles were precipitated using PEG and added to the cell lysate. The cells were lysed using a salt active nuclease (SRE0015, Sigma) and centrifuged to form a pellet. The supernatant containing the AAV particles was then loaded onto an iodixanol gradient and centrifuged for 2.25 h at 59.000 RPM (70 Ti rotor) in an ultracentrifuge. 4-5 mL of liquid was collected from the clear 40% layer containing the viral particles and washed and concentrated using Amicon filters to the desired volume. AAV titers were estimated using qPCR and were later aliquoted and kept at −80°C for long-term storage.

### Viral injection and cranial window implantations

Viral injections and cranial window implantation were performed as previously described ^28,63^. Briefly, mice were anesthetized with isoflurane and fixed in a stereotaxic apparatus (Kopf Instruments, model 942) and body temperature was maintained using a heating pad set at 37°C. Ophthalmic ointment (Duratears) was applied to the eyes and local anesthesia (2% Lidocaine) was injected under the scalp and the skin was cleaned with 70% ethanol and betadine. Craniotomies were made above the left visual cortex (centered 2.7 mm posterior and 2.5 mm lateral to bregma). For imaging experiments, a 4mm craniotomy was made above the visual cortex to allow for window implantation. 2-3 injections of 300nL of virus were made using a beveled glass micropipette at a depth of ∼300 μm to target layer 2/3 of the cortex at a rate of 65 nl/min with a microsyringe pump (UMP3T-2, WPI). For imaging experiments, a cover glass and custom-made 3D printed head post painted black was glued to the skull using cyanoacrylate glue (Krazy glue). For all other experiments, the skin of the animal was sealed with VetBond (3M). Following surgery, animals were administered with analgesic (0.1 mg/kg of buprenorphine and 5 mg/kg of Carprofen). After initial recovery on a heating pad (RWD Life Science), mice were returned to their homecage and monitored for post-op care. We injected the following AAV titers in the *Npas4* knockout experiments: the AAVs of the pAAV-CW3SL-mRuby3-2A-Cre-WPRE and pAAV-CW3SL-mRuby3-2A-dCre-WPRE constructs were injected for *Npas4* knockout calibration experiments (**Supplemental Figure 4B-D**) at concentrations of 2.06 E+14 and 1.84 E+14 viral units per microliter, respectively, and at ten-fold dilutions for the *Npas4* knockout *ex vivo* electrophysiology experiments (**Figure 4**) and *in vivo* calcium imaging experiments (**Figure 5**).

### Perfusions and immunohistochemistry

#### Perfusion

Animals were anesthetized with pental and were transcardially perfused with 5 mL of ice-cold 1x PBS and subsequently 15 mL of ice-cold fixative (4% PFA in PBS). Brains were then dissected, post-fixed (in 4% PFA) for 1 hour on a rotator at 4°C and washed three times in cold PBS. Brains were placed overnight in 30% sucrose at 4°C and subsequently embedded and frozen in OCT compound (TissueTek, #4583). Coronal sections of tissue were cut using a cryostat (Leica CM1950) and mounted on SuperFrost Plus glass slides (Fisher Scientific).

#### Immunohistochemistry

Coronal sections (10 µm thick) of AAV infected visual cortex tissue were cut using a Leica CM1950 cryostat and mounted on SuperFrost Plus glass slides (Fisher Scientific). The slices were washed in PBS and incubated for 1 hour in a blocking solution (1x PBS, 10% NGS, 0.5% Triton-X). The blocking buffer was then replaced with the Rabbit-anti-Npas4 antibody (Activity Signalling, Lot #NP41-1) diluted in the blocking buffer (1:1000) and left at 4°C for 60 hours. The slides were then washed in PBS and incubated for 1 hour at room temperature with Goat-anti-rabbit Alexa 488 (Molecular Probes, A-11039) diluted 1:1000 in a blocking buffer (1x PBS, 10% NGS, 0.3% Triton-X) with a nuclear stain (Hoechst 33342). Slides were mounted with Fluoromount-G (SouthernBiotech, 0100-01) and imaged on a confocal microscope (Zeiss LSM800).

### RNA extraction

Visual cortices were dissected from adult mice (∼P66) along the day either in the dark or following light-exposure. Animals that were not light-stimulated were enucleated in the dark with night-vision goggles prior to dissecting the tissue. Dissected tissues were snap-frozen and stored at −80C until further processing. RNA was homogenized and extracted in a 2mL Dounce homogenizer containing TRIzol Reagent (Thermo Fisher Scientific) using a standard TRIzol Reagent-based RNA extraction protocol. RNA was then further cleaned using Agencourt RNAClean XP beads, and concentration was determined using NanoDrop One Spectrophotometer (Thermo Fisher Scientific). RNA quality was validated using the 2200 TapeStation (Agilent technologies).

### RNA sequencing and bioinformatic analysis

Bulk RNA-seq libraries were prepared at the Crown Genomics Institute of the Nancy and Stephen Grand Israel National Center for Personalized Medicine, Weizmann Institute of Science (INCPM). Libraries were prepared based on the Mars-Seq protocol ^90^, which allows for sequencing of the 3’ ends of RNA transcripts. Prepared libraries were sequenced on the NovaSeq 6000 using the Illumina kit NovaSeq 6000 SP Reagent Kit v1.5 (100 cycles).

Processing of the raw read files (fastq) was performed via UTAP: User-friendly Transcriptome Analysis Pipeline (v1.0.9 ^91^). Briefly, reads were trimmed using cutadapt (v.1.15 ^92^) and mapped to the mouse genome (mm10) using STAR (v.2.5.2b ^93^). UMIs (unique molecular identifiers) were counted using HTSeq-count (v.0.9.1 ^94^).

### Home Cage Running experiments

Adult wild type animals were individually housed in standard circadian cabinets (Actimetrics) with plastic running wheels to monitor running of individual animals. Animals were allowed at least 7 days of acclimation to running wheels prior to experimental start. Locomotion activity (number of wheel rotations per minute) was monitored using ClockLab data collection software (Actimetrics). On the final day of the experiment, light was not presented but no other changes to the animal’s environment were made. Running wheel activity was analyzed using custom written matlab scripts. Running was binned into 1 hour bins, and was normalized to the maximum rotations per hour across the total recording session for each individual animal recorded. Average running of all mice on DLD days was compared with their activity on DDD days.

### Fluorescent *in situ* hybridization (FISH)

For RNAscope FISH in acute brain slices, brain tissues were freshly dissected and frozen in OCT compound (TissueTek, #4583). Coronal sections (15 µm thick) of the visual cortex were cut using a Leica CM1950 cryostat and mounted on SuperFrost Plus glass slides (Fisher Scientific). FISH was performed following the RNAscope Fluorescent Multiplex Assay protocol with pretreatment III according to the manufacturer’s instructions. The probes against *Npas4*, *Slc17a* and *Fos* were used to measure gene expression in excitatory neurons. For measuring gene expression in VIP INs probes were used against *Igf1* and *Vip*. The ACD 3-plex negative probe control set was used to determine the background in the respective channels. RNAscope FISH images were analyzed in Matlab with custom-written scripts as previously described ^28^ to quantify the number of *Npas4, Fos* and *Igf1* puncta present per nucleus; data presented in violin plots (**Figure 1E**, **Supplemental Figure 1G,H**) are based on background-subtracted of expression values of *Npas4* and *Fos* in single cells.

### Patch clamp electrophysiology in acute visual cortex slices

P60-80 mice were transcardially perfused with ice-cold sucrose dissection media (26 mM NaHCO_3_, 1.25 mM NaH_2_PO_4_, 2.5 mM KCl, 10 mM MgSO_4_, 11 mM glucose, 0.5 mM CaCl_2_, 234 mM sucrose; 340 mOsm). Brains were then dissected and sliced, while being kept in ice-cold sucrose dissection media, into coronal sections (300 μm thick) containing the primary visual cortex using a Leica VT1200S vibratome. Slices were incubated in high osmotic concentrated artificial cerebrospinal fluid (28.08 mM NaHCO_3_, 1.35 mM NaH_2_PO_4_, 132.84 mM NaCl, 3.24 mM KCl, 1.08 mM MgCl_2_, 11.88 mM glucose, 2.16 mM CaCl_2_; 320 mOsm) at 32°C for 30 minutes immediately after slicing. Then, slices were incubated in normal osmotic concentrated artificial cerebrospinal fluid (26 mM NaHCO_3_, 1.25 mM NaH2PO_4_, 123 mM NaCl, 3 mM KCl, 1 mM MgCl_2_, 11 mM glucose, 2 mM CaCl_2_; 300 mOsm) at 32°C for 30 minutes and subsequently at room temperature. All solutions were saturated with 95%-O2/5%-CO_2_, and slices were used within 6 hours of preparation. Whole-cell patch-clamp recordings were performed in aCSF at 32°C from neurons in the visual cortex that were identified under fluorescent and DIC optics. Recording pipettes were pulled from borosilicate glass capillary tubing with filaments (OD 1.50 mm, ID 0.86 mm, length 10 cm) using a P-1000 micropipette puller (Sutter Instruments) and yielded tips of 2–5 MΩ resistance. Recordings were sampled at 20 kHz and filtered at 3 kHz.

All mPSC experiments were recorded with pipettes filled with an internal solution containing: 135 mM caesium methanesulfonate, 5 mM TEA-Cl, 10 mM HEPES, 4 mM Mg-ATP, 0.3 mM Na-GTP, 0.3 mM EGTA, and 3 mM QX-314-Cl. Osmolarity and pH were adjusted to 310 mOsm and 7.3 pH with sucrose and CsOH, respectively. mEPSCs were isolated from mIPSCs by exposing neurons to 0.5 μM tetrodotoxin (TTX) and holding them intermittently at −70 mV or 0 mV, respectively, as previously described ^74^. For measuring the intrinsic properties the following internal solution was used: 135 mM k-gluconate, 4mM KCl, 10mM HEPES, 10mM Pcreatine, 4mM Mg-ATP, 4mM GTP-Na and 2mM Na2-ATP. Intrinsic properties were calculated by giving 500ms long current steps (5pA); no TTX was added to the extracellular solution in these experiments. Data were acquired via Clampex10 using a Multiclamp 700B amplifier, and digitized with an Axon Digidata 1550B data acquisition board (Axon Instruments.

We were able to identify L2/3 pyramidal neurons by their morphology under bright field and their large capacitance (patched neurons with capacitance less than 60 pico-farad (pF) were excluded from the dataset). Visualization of pyramidal neurons was conducted with an Olympus U-CMAD4 T7 microscope (Olympus, Tokyo, JPN) together with a LUMPlan FI/L 40x water immersion objective mounted on a TMC Micro-g air table (TMC, Massachusetts, USA) with a Hamamatsu Orca – FLASH 3.0.

The following numbers of neurons were recorded from each mouse: **Figure 2C**: 4 mice per condition; ZT0 - 9/7/14/7 cells per mouse; ZT1 - 11/8/9/11 cells per mouse; ZT4 - 6/12/13/6 cells per mouse; ZT8 - 8/13/10/12 cells per mouse; ZT12 - 8/11/11/10 cells per mouse; ZT1 Dark - 6/6/4/5 cells per mouse; ZT12 Dark - 12/6/5/15 cells per mouse. **Figure 4B,C**: 4-5 mice per condition; 7 days Dark Control - 9/10/10/8 cells per mouse; 7 days Dark followed + 12 hours Light Control - 12/12/11/11; 7 days Dark cKO - 9/8/5/9 cells per mouse; 7 days Dark + 12 hour Light cKO - 9/6/8/8/11. **Figure 4E,F**: 4 mice per condition; ZT 0 Control - 8/7/9/10 cells per mouse; ZT 0 cKO - 7/10/11/9 cells per mouse; ZT1 Control - 8/6/10/6 cells per mouse; ZT1 cKO - 6/5/6/4 cells per mouse; ZT 12 Control - 5/4/11/5 cells per mouse; ZT 12 cKO - 7/10/9/8 cells per mouse. **Supplemental Figure 2E,F**: 4 mice per condition; ZT 0 - 5/6/13/11 cells per condition; ZT 1 - 5/5/11/9 cells per condition; ZT 12 - 8/7/9/5 cells per condition.

### Longitudinal *In vivo* 2-photon imaging

#### In vivo 2-photon calcium imaging

Imaging was performed 2-5 weeks following AAV injections using a two-photon microscope equipped with a 12 kHz resonant-galvo scanhead (Bergamo microscope, ThorLabs) while the animals were head-fixed but not body-restrained under the microscope. Illumination was provided by a Ti:Sapphire laser (MaiTai DeepSee, Spectra Physics) tuned to a wavelength of 930 nm. Images were acquired in three planes 30 um apart simultaneously using a fast piezo motor (Thorlabs). Frame size was 512 x 512 pixels, allowing for a frame acquisition rate of 11.4 Hz per plane. For cKO experiments, images of red fluorescence were acquired following functional imaging using 1040nm illumination. FOVs were matched across recording sessions according to blood vessel pattern and fine structural details. To limit stress of animals due to repeated and extended times of head fixation, animals were recorded up to twice per day with a minimum of 12 hours in between initiation of recording sessions.

#### Visual stimulation

Visual stimuli were generated using Matlab Psychophysics toolbox (http://psychtoolbox.org/) and presented on a gamma corrected LCD screen. The screen was positioned 20 cm from the contralateral eye of the recorded hemisphere. Stimuli were presented for 1 second with an interstimulus interval of 4 seconds during which a gray screen of mean luminance was presented. Visual stimuli consisted of full field sinusoidal drifting bar gratings with a temporal frequency of 2 Hz. Varying contrasts, spatial frequencies and directions (equal spacing of 30 degrees) were presented in a pseudorandom order. 14-15 repetitions were presented per stimulus during each recording session. During recordings where ongoing activity, and not visually evoked activity, was recorded a static gray screen of mean luminance was displayed to the animal.

#### FOV mapping

Field of views (FOVs) were mapped in order to determine the location of V1 and best imaging area for longitudinal recordings per animal. Receptive fields were mapped by presenting patches of drifting sinusoidal gratings spaced on a 3 by 4 grid of the monitor. Stimuli were presented for 1 second at 0 degrees, 100% contrast and 0.04 cpd with an interstimulus interval of 4 seconds. FOVs were selected based on receptive field mapping and percentage of visually responsive neurons in the area.

#### Pupillometry and facial movements

During imaging, the mouse’s ipsilateral (unstimulated) eye and whisker pad was illuminated with an IR-light source (M940L3, Thorlabs) and imaged using a CMOS camera at 33 Hz. Pupillometry data was synchronized using flashing of the IR camera.

### Data analyses

#### RNA-seq

Following the run of UTAP, all subsequent gene expression analyses were performed in R. Normalization of expression levels and differential expression testing were performed using the DESeq2 package ^95^. Genes were flagged as expressed if they passed a minimal expression threshold of 10 reads in at least 3 of the 15 collected time points, resulting in 10,805 genes. Next, genes were flagged as regulated between time points (compared with the ZT0/ZT11.5 timepoints) if they answered the following criteria: 1) P value <0.01, 2) absolute mean fold change was of 1.75-fold or greater, 3) absolute fold changes of 1.75 or higher in at least 3 of 4 biological replicates (the Dark ZT1, Dark ZT13 and Light ZT18 time points had 3 biological replicates and not 4- in these cases, a fold change of 1.75 or higher was required in at least 2 of 3 biological replicates), and 4) the mean expression value in one of the two compared samples must be above the 15th percentile expression threshold. Using this threshold, 146 genes were found to be regulated.

Gene expression plots show the average and standard error of the mean of 4 biological replicates (and 3 biological replicates in the case of Dark ZT1, Dark ZT13 and Light ZT18 time points), all containing a mix of both male and female mice.

#### Gene ontology (GO)

To further explore the functional roles of the genes in the different groups of regulated genes (e.g. sensory-regulated genes, dark-regulated genes), we performed gene ontology (GO) analysis using the Gene Ontology Resource (http://geneontology.org/). Presented results show representation in the GO categories of biological processes, molecular functions and cellular components. Only GO terms with FDR-corrected P-values <0.05 were included in results.

#### Electrophysiological recordings

All mPSC events were automatically detected and characterized as previously described ^28^ with custom-written MATLAB scripts using the functions abfload and iPeak from MATLAB Central File Exchange. The charge was calculated as the area under the curve of each single mPSC event (i.e., only the areas under curves of detected events were used for the calculation of the charge). The starting and ending points of an event were determined as the nearest time points before and after a peak where the value of the measured current was lower than 10% of the value of the event’s amplitude. The E/I-ratio was calculated based on the average charge (C) of mPSCs, as follows:

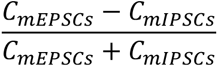

The average mPSC charge was calculated as the average sum of current along one second of recording (as 1 Coulomb = 1 Ampere ⋅ 1 Second). All data were analyzed blind to genotype or experimental conditions. Statistical tests were performed using Graphpad Prism. Mann-Whitney test was used for paired comparisons. In case of multiple comparisons, post-hoc Dunnett’s T3 multiple comparisons test was performed following a Brown-Forsythe ANOVA test or Kruskall - Wallis.

#### Pupil diameter and facial movement

Analyses of pupil diameter and facial movement were performed as previously described ^63^. Videos were analyzed to extract pupil diameter and facial movement (i.e. whisking) using a custom written LabView GUI. Pupil diameter analysis-Frames were filtered using a median filter and thresholded to low IR light reflectance areas. The resulting areas were filtered based on circularity and size until only the region corresponding to the pupil remained. An ellipse was then fitted to this region by setting its major and minor axes equal to the longer and shorter lines of symmetry of the bounding rectangle. The pupil diameter was estimated as the geometric mean of the major and minor axes. Raw pupil signal was smoothed using a Savitzky–Golay filter with a first order polynomial. Facial energy measurement - The squared difference in intensity between the pixels of each pair of consecutive frames (i.e. mean squared error between frames) was calculated. Raw whisking signal was smoothed using an Savitzky–Golay filter with a second order polynomial, and whisking episodes were identified as high-amplitude changes in the signal crossing the threshold, with at least 300 ms in duration. Pupil diameter and facial energy traces were synchronized with calcium F traces using custom written Matlab scripts.

#### In vivo 2-photon calcium imaging data

Raw movies were analyzed using Suite2p software suite ^96^, where movies were registered, ROIs were detected, and fluorescent traces (F) and deconvolved signals were computed. Cells were matched across recording sessions using RoiMatchPub (https://github.com/ransona/ROIMatchPub) or CellReg algorithms (https://github.com/zivlab/CellReg). Following signal extraction and cell matching, all recordings were analyzed using custom software written in house in MATLAB (The MathWorks). F signals were neuropil corrected and aligned to timing of visual stimulation and arousal state data (i.e. pupil size, whisking) using frame time signals collected in Clampex (Molecular Devices) during recording sessions.

#### Ongoing neural activity

Deconvolved signal was used in order to estimate the amount of ongoing activity of each cell at each recording session. The deconvolved signal of each cell was summed across ∼15 minutes of ongoing activity recording. In order to compare matched cells across the day, each cell’s activity was normalized to its activity at ZT0. To minimize single animal driven effects, cells were randomly subsampled to result in an equal number of cells per animal at each timepoint.

#### Visually evoked neural activity

Visually evoked responses were calculated as the mean ΔF/F during visual stimulus presentation:

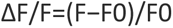

Where F0 was calculated as the mean fluorescence during the 1 second preceding the visual stimulus. To determine if a cell was visually responsive, a one-way non-parametric ANOVA (Kruskall-Wallis) was computed comparing the evoked responses to baseline fluorescence. An additional non-parametric test was performed between the activity during the interstimulus interval and during presentation of the preferred visual stimulus (Wilcoxon Rank Sum test, p < 0.05). Cell subgroups were classified based on how often the matched cell was visually responsive, such that ‘Ever’ visually responsive cells were statistically responsive during at least of the recorded sessions while ‘Stable’ visually responsive cells were visually responsive during all recorded sessions.

The preferred stimulus type (direction, spatial frequency, or contrast) was defined as the stimulus that elicited the largest response across the tested parameters.

Orientation selectivity was calculated as previously described (Mazurek et al., 2014) using a calculation of the global orientation selectivity index (gOSI):

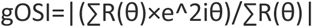

Where R(θk) is the response to angle θk. To calculate the population average, responses of each cell were normalized to their maximum response and artificially centered to 0°. Normalized responses of cells were averaged and fitted using a double Gaussian:

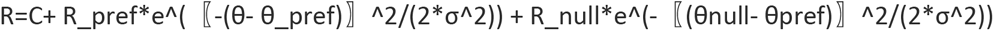

#### Immuno-labeling fluorescent Image analyses

NPAS4 protein levels were measured with a custom-written Image J script. Per cell, the relative background noise was calculated and subtracted from the raw fluorescent values. If the cell’s resulting value was negative (i.e. the background noise was higher than the raw value) it was substituted by 0. The same analysis was performed to identify the mRuby positive cells (AAV infected).

### Statistics

The number of experimental recordings and animals used in each experiment is indicated in the figure legends or in the respective supplemental tables. Statistical tests were performed using MATLAB, p values and statistical tests used are indicated in the figure legends.

---

**Supplemental Figure 1.**
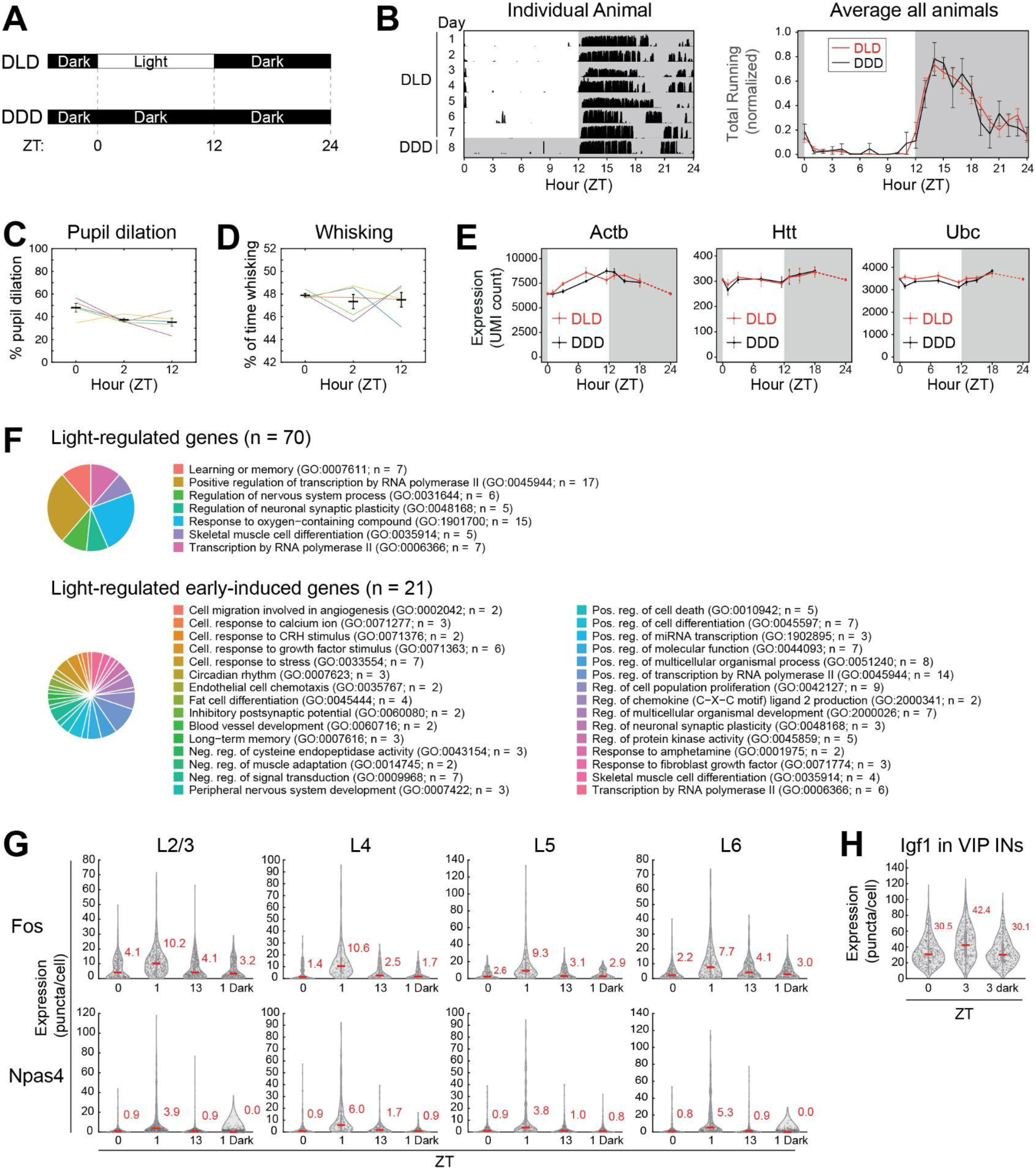
Additional data and analyses related to daily light-induced changes in gene expression in the visual cortex. (Related to Figure 1) **A, B** Running activity of mice is not disrupted by the lack of light on a single day (i.e. in DDD cohorts). **A** Experimental strategy. Adult mice were raised and housed under standard housing conditions in a standard 12/12 hour dark-light cycle. On the day of the experiments, light appeared as usual for mice during the Dark-Light-Dark (DLD) days but not on the Dark-Dark-Dark (DDD) day when the animals remained in the dark. **B** (Right) Running pattern of a representative mouse across the seven days before the experiment (i.e. on the DLD days under standard 12/12 hour dark-light conditions) and on the day when the light did not appear (i.e. on the DDD day); black lines indicate running, shaded areas indicate times of darkness. (Left) Average running of all mice on DLD days (red) and the DDD day (black); shaded area indicates times of expected darkness. No significant differences were observed between the running patterns of the animals on DLD and DDD days. **C, D** Arousal state as indicated by pupil dilation (**C**) and whisking (**D**) does not change significantly across the day’s light period. Data of individual animals is shown in color, while the averages across all animals are in black. **C** Pupillometry. Average percent pupil size of longitudinally recorded mice (n = 5). **D** Whisking. Percent of time spent whisking of longitudinally recorded mice (n = 5). **E** The expression of housekeeping genes is not altered across the day and is not altered in response to the daily appearance of light (i.e. in DLD cohorts) or its lack on one day (i.e. in DDD cohorts). **F** GO-term analyses of light-regulated genes. Top: Biological processes associated with all light-regulated genes (n=70 genes). Bottom: Biological processes associated with early-induced light-regulated genes (n=21 genes). **G** Layer-specific quantification of RNAscope smFISH puncta for *Fos* (top) and *Npas4* (bottom) in *Slc17a7*-positive visual cortex excitatory neurons at ZT0/1/13 in DLD mice or ZT1 in DDD mice (= ZT1 Dark) (red line & number = median). **H** Quantification of RNAscope smFISH puncta for *Igf1* in *Vip*-positive visual cortex GABAergic interneurons at ZT0/3 in DLD mice or ZT3 in DDD mice (red line & number = median; see Supplemental Figure 1G for layer-specific quantification). (Error bars represent SEM in all data panels)

**Supplemental Figure 2.**
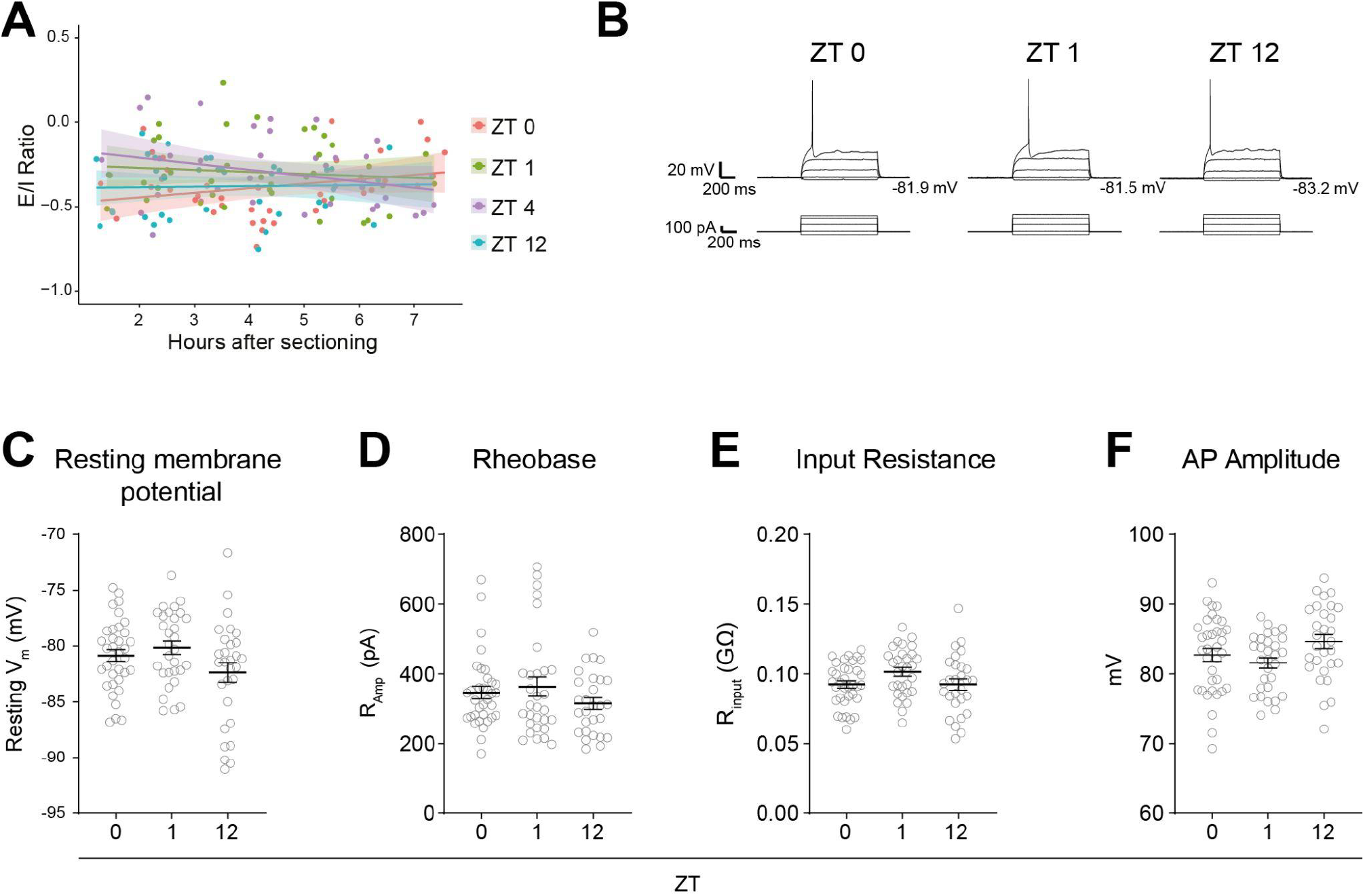
The intrinsic properties of L2/3 PYR neurons in the visual cortex are not altered by the daily appearance of light. (Related to Figure 2) **A** Scatter plot of the E/I ratio of L2/3 PYR neurons in acute visual cortex slices of DLD mice sacrificed at ZT 0, 1, 4 and 12 relative to the time after preparation of the slices. No significant correlation between E/I-ratio and time of recording after slice preparation was observed. **B** Representative traces of patch clamp recordings in current clamp mode in L2/3 PYR neurons in acute visual cortex slices of DLD mice sacrificed at ZT 0, 1 and 12. The traces denote the neurons’ voltage response (top) relative to the injected current (bottom). **C-F** Quantified data of Resting membrane potential, Rheobase, Input resistance and Action potential amplitude in L2/3 PYR neurons in acute visual cortex slices of DLD mice sacrificed at ZT 0, 1 and 12 (ZT 0 n = 35 cells from 4 mice; ZT 1 n = 30 cells from 4 mice; ZT 12 n = 29 cells from 4 mice; Statistics Kruskal-Wallis test with post-hoc Dunnett’s T3 multiple comparisons test: no statistically significant differences were observed). (Error bars represent SEM in all data panels; see **Supplemental Data Table 2** for descriptive statistics)

**Supplemental Figure 3.**
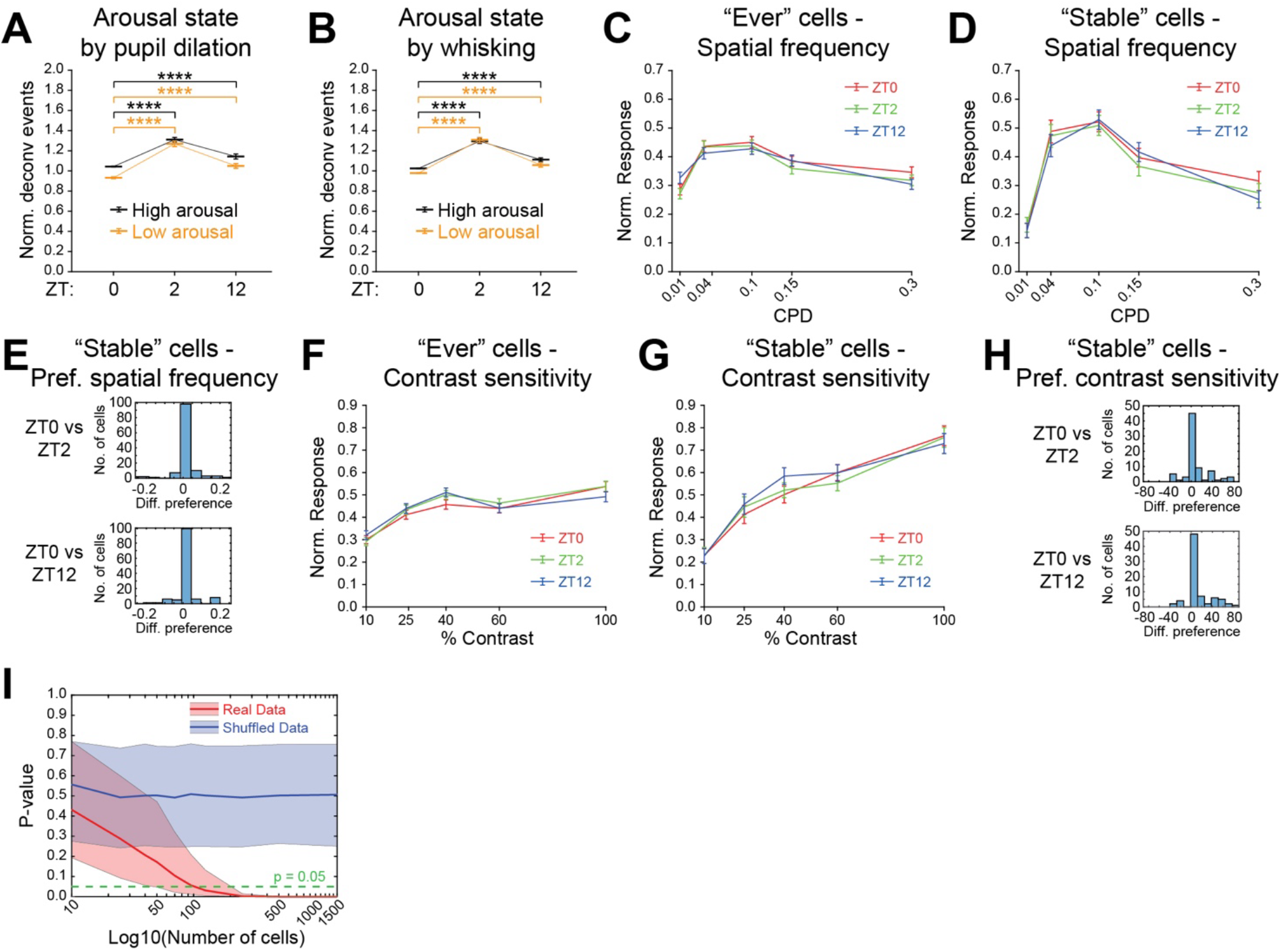
Additional data and analyses related to daily light-induced changes in neural activity and tuning properties of L2/3 excitatory neurons in the visual cortex. (Related to Figure 3) **A, B** Daily changes in ongoing activity are not driven by differences in arousal state. Normalized activity of neurons at low (yellow) and high (black) arousal state, as measured by normalized pupil size (**A**) or whisking events (**B**) (n = 1275 cells from 5 mice; Statistics Wilcoxon Sign Rank test **** p<0.0001) **C-E** Spatial frequency tuning and preference are not altered during the day’s light phase in DLD mice (ZT0 = red, ZT2 = green, ZT12 = blue). Normalized population averages of ‘Ever’ (**C**) and ‘Stable’ (**D**) visually responsive neurons (‘Ever’, n = 418 cells; ‘Stable’, n = 127 cells). **E** Histograms of difference in spatial frequency preference of ‘Stable’ neurons between ZT0 and ZT2 (top) or ZT12 (bottom). **F-H** Contrast sensitivity tuning and preference are not altered during the day’s light phase in DLD mice (ZT0 = red, ZT2 = green, ZT12 = blue). Normalized population averages of ‘Ever’ (**F**) and ‘Stable’ (**G**) visually responsive cells (‘Ever’ n = 340 cells; ‘Stable’ n = 77 cells). **H** Histograms of difference in contrast sensitivity preference of ‘Stable’ neurons between ZT0 and ZT2 (top) or ZT12 (bottom). **I** Average p-value of 5000 simulations of comparisons between increasing numbers of randomly sub-selected matched neurons at ZT0 and ZT2 (solid lines = median p-values; shaded areas = 25th/75th percentile; shuffled data as negative control). (Error bars represent SEM in all data panels; see **Supplemental Data Table 3** for descriptive statistics)

**Supplemental Figure 4.**
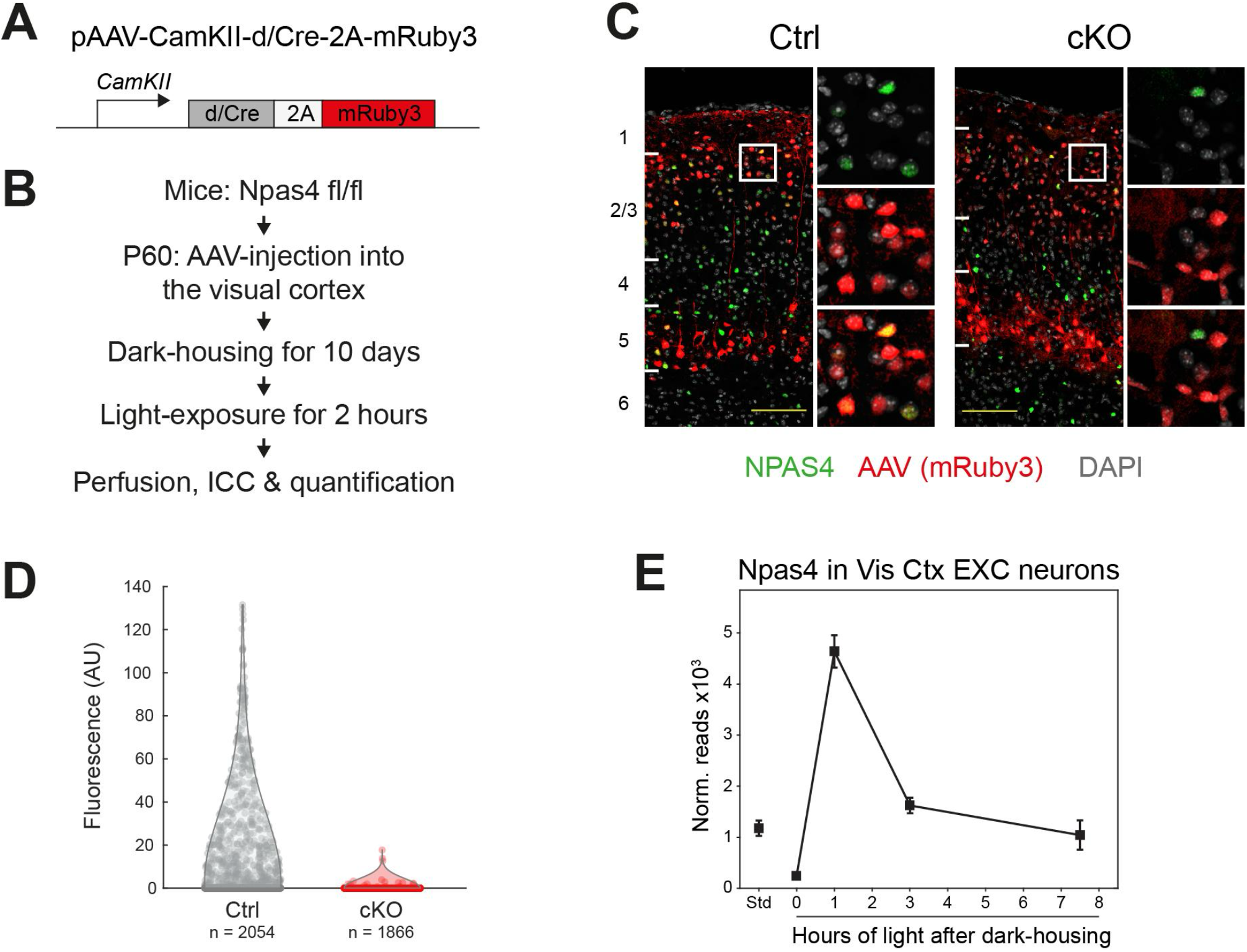
Validation of an AAV-based strategy for Cre-conditional knock-out of *Npas4* in excitatory neurons in the adult visual cortex. (Related to Figures 4 and 5) **A** Design of the AAV construct for expression of Cre-recombinase (dCre as negative control) and the red-fluorescent protein mRuby3 in cortical excitatory neurons via a CamKIIa promoter (W3SL promoter ^97^). **B** Experimental strategy to test AAV-mediated cKO of NPAS4 in the visual cortex of adult *Npas4* fl/fl mice. **C** Representative images of visual cortex sections of Npas4 fl/fl mice injected with either the control (dCre) or cKO (Cre) AAV and immuno-labeled with antibodies against NPAS4 (green = anti-NPAS4; red = AAV-infected neurons; scale-bar = 100 microns; numbers on the left indicate cortical layers). **D** Quantification of NPAS4 fluorescence in AAV infected (=mRuby3-positive) neurons in visual cortex sections of Npas4 fl/fl mice injected with either the control (dCre) or cKO (Cre) AAV and immuno-labeled with antibodies against NPAS4 (Control: n = 2054 mRuby3-positive cells from 3 mice; cKO: n = 1866 mRuby3-positive cells from 3 mice). **E** RiboTag-seq data of *Npas4* in visual cortex excitatory neurons of adult mice that were either housed in a standard 12/12 hours dark-light cycle (Std) or dark-housed for two weeks and then exposed to 0, 1, 3 or 7.5 hours of light (data taken from ^37^; error bars represent SEM).

**Supplemental Figure 5.**
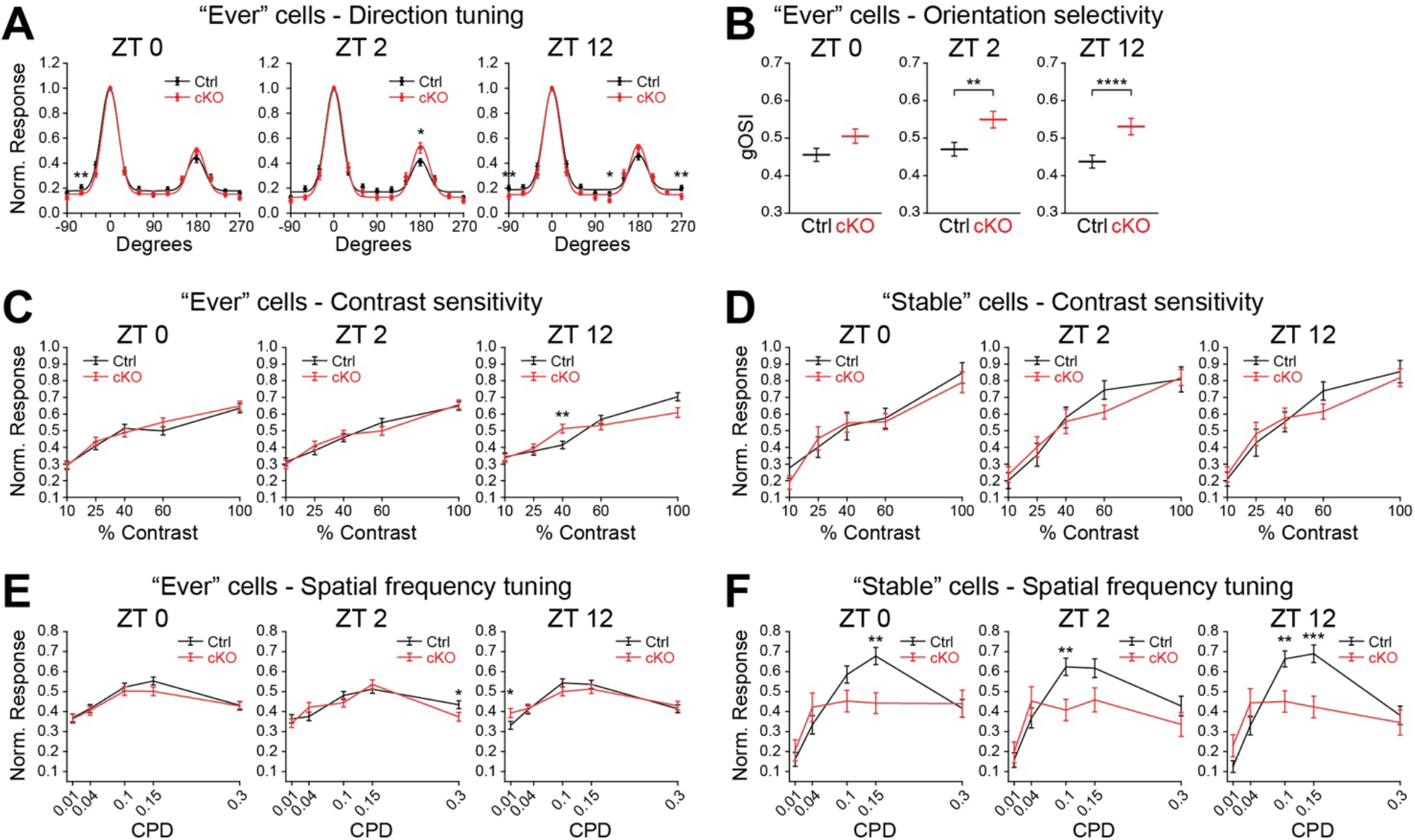
Additional analyses of the tuning properties in visual cortex L2/3 PYR neurons upon cKO of Npas4. (Related to Figure 5) **A** Direction tuning of Control (black) or cKO (red) ‘Ever’ visually responsive neurons. Normalized and double-gaussian-fitted population averages at ZT0 (left), ZT2 (middle) or ZT12 (right) of visually responsive neurons (lines) and normalized average responses (diamonds) are shown (Wilcoxon Sign Rank test * p<0.05, ** p<0.01). **B** Global orientation selectivity indices (gOSI) of ‘Ever’ visually responsive neurons (Wilcoxon Signed-Rank test, ** p<0.01, **** p<0.0001). **C, D** Contrast sensitivity tuning of Control (black) or cKO (red) ‘Ever’ (**C**) and ‘Stable’ (**D**) visually responsive neurons (‘Ever’ neurons: Ctrl n = 221 cells, cKO n = 191 cells; ‘Stable’ neurons: Ctrl n = 38 cells, cKO n = 42 cells; Statistics Wilcoxon Rank Sum (**C)** or Wilcoxon Sign Rank (**D**) test * p<0.05, ** p<0.01). **E, F** Spatial frequency tuning of Control (black) or cKO (red) ‘Ever’ (**E**) and ‘Stable’ (**F**) visually responsive neurons (‘Ever’ neurons: Ctrl n = 387 cells, cKO = 330 cells; ‘Stable’ neurons: Ctrl n = 113 cells, cKO = 92 cells; Statistics Wilcoxon Rank Sum (**E)** or Wilcoxon Sign Rank (**F**) test * p<0.05, ** p<0.01, *** p<0.001). (Error bars represent SEM in all data panels; see **Supplemental Data Table 3** for descriptive statistics)

## References

1. Citri, A., and Malenka, R.C. (2007). Synaptic Plasticity: Multiple Forms, Functions, and Mechanisms. Neuropsychopharmacology 33, 18–41.

2. Morrison, A., Aertsen, A., and Diesmann, M. (2007). Spike-timing-dependent plasticity in balanced random networks. Neural Comput. 19, 1437–1467.

3. Litwin-Kumar, A., and Doiron, B. (2014). Formation and maintenance of neuronal assemblies through synaptic plasticity. Nat. Commun. 5, 5319.

4. Zenke, F., Hennequin, G., and Gerstner, W. (2013). Synaptic plasticity in neural networks needs homeostasis with a fast rate detector. PLoS Comput. Biol. 9, e1003330.

5. Magloire, V., Mercier, M.S., Kullmann, D.M., and Pavlov, I. (2019). GABAergic interneurons in seizures: Investigating causality with optogenetics. Neuroscientist 25, 344–358.

6. Douglas, R.J., Koch, C., Mahowald, M., Martin, K.A., and Suarez, H.H. (1995). Recurrent excitation in neocortical circuits. Science 269, 981–985.

7. Chen, J.-Y., Lonjers, P., Lee, C., Chistiakova, M., Volgushev, M., and Bazhenov, M. (2013). Heterosynaptic plasticity prevents runaway synaptic dynamics. J. Neurosci. 33, 15915–15929.

8. Abbott, L.F., and Nelson, S.B. (2000). Synaptic plasticity: taming the beast. Nat. Neurosci. 3 Suppl, 1178–1183.

9. Marder, E., and Prinz, A.A. (2003). Current compensation in neuronal homeostasis. Neuron 37, 2–4.

10. Turrigiano, G.G. (2017). The dialectic of Hebb and homeostasis. Philos. Trans. R. Soc. Lond. B Biol. Sci. 372. 10.1098/rstb.2016.0258.

11. Keck, T., Toyoizumi, T., Chen, L., Doiron, B., Feldman, D.E., Fox, K., Gerstner, W., Haydon, P.G., Hübener, M., Lee, H.-K., et al. (2017). Integrating Hebbian and homeostatic plasticity: the current state of the field and future research directions. Philos. Trans. R. Soc. Lond. B Biol. Sci. 372. 10.1098/rstb.2016.0158.

12. Lee, H.-K., and Kirkwood, A. (2019). Mechanisms of Homeostatic Synaptic Plasticity in vivo. Front. Cell. Neurosci. 13, 520.

13. Slomowitz, E., Styr, B., Vertkin, I., Milshtein-Parush, H., Nelken, I., Slutsky, M., and Slutsky, I. (2015). Interplay between population firing stability and single neuron dynamics in hippocampal networks. Elife 4, 126–121.

14. Keck, T., Keller, G.B., Jacobsen, R.I., Eysel, U.T., Bonhoeffer, T., and Hübener, M. (2013). Synaptic scaling and homeostatic plasticity in the mouse visual cortex in vivo. Neuron 80, 327–334.

15. Hengen, K.B., Lambo, M.E., Van Hooser, S.D., Katz, D.B., and Turrigiano, G.G. (2013). Firing rate homeostasis in visual cortex of freely behaving rodents. Neuron 80, 335–342.

16. Barnes, S.J., Sammons, R.P., Jacobsen, R.I., Mackie, J., Keller, G.B., and Keck, T. (2015). Subnetwork-Specific Homeostatic Plasticity in Mouse Visual Cortex In Vivo. Neuron 86, 1290–1303.

17. Torrado Pacheco, A., Bottorff, J., Gao, Y., and Turrigiano, G.G. (2021). Sleep Promotes Downward Firing Rate Homeostasis. Neuron 109, 530–544.e6.

18. Radulescu, C.I., Doostdar, N., Zabouri, N., Melgosa-Ecenarro, L., Wang, X., Sadeh, S., Pavlidi, P., Airey, J., Kopanitsa, M., Clopath, C., et al. (2023). Age-related dysregulation of homeostatic control in neuronal microcircuits. Nat. Neurosci. 26, 2158–2170.

19. Pecoraro-Bisogni, F., Lignani, G., Contestabile, A., Castroflorio, E., Pozzi, D., Rocchi, A., Prestigio, C., Orlando, M., Valente, P., Massacesi, M., et al. (2018). REST-Dependent Presynaptic Homeostasis Induced by Chronic Neuronal Hyperactivity. Mol. Neurobiol. 55, 4959–4972.

20. Howard, M.A., Rubenstein, J.L.R., and Baraban, S.C. (2014). Bidirectional homeostatic plasticity induced by interneuron cell death and transplantation in vivo. Proc. Natl. Acad. Sci. U. S. A. 111, 492–497.

21. Stellwagen, D., and Malenka, R.C. (2006). Synaptic scaling mediated by glial TNF-alpha. Nature 440, 1054–1059.

22. Seeburg, D.P., Feliu-Mojer, M., Gaiottino, J., Pak, D.T.S., and Sheng, M. (2008). Critical role of CDK5 and Polo-like kinase 2 in homeostatic synaptic plasticity during elevated activity. Neuron 58, 571–583.

23. Davis, G.W. (2006). Homeostatic control of neural activity: from phenomenology to molecular design. Annu. Rev. Neurosci. 29, 307–323.

24. Shepherd, J.D., Rumbaugh, G., Wu, J., Chowdhury, S., Plath, N., Kuhl, D., Huganir, R.L., and Worley, P.F. (2006). Arc/Arg3.1 mediates homeostatic synaptic scaling of AMPA receptors. Neuron 52, 475–484.

25. Rutherford, L.C., Nelson, S.B., and Turrigiano, G.G. (1998). BDNF has opposite effects on the quantal amplitude of pyramidal neuron and interneuron excitatory synapses. Neuron 21, 521–530.

26. Desai, N.S., Rutherford, L.C., and Turrigiano, G.G. (1999). Plasticity in the intrinsic excitability of cortical pyramidal neurons. Nat. Neurosci. 2, 515–520.

27. Barnes, S.J., Keller, G.B., and Keck, T. (2022). Homeostatic regulation through strengthening of neuronal network-correlated synaptic inputs. Elife 11. 10.7554/eLife.81958.

28. Roethler, O., Zohar, E., Cohen-Kashi Malina, K., Bitan, L., Gabel, H.W., and Spiegel, I. (2023). Single genomic enhancers drive experience-dependent GABAergic plasticity to maintain sensory processing in the adult cortex. Neuron 111, 2693–2708.e8.

29. Favuzzi, E., Marques-Smith, A., Deogracias, R., Winterflood, C.M., Sánchez-Aguilera, A., Mantoan, L., Maeso, P., Fernandes, C., Ewers, H., and Rico, B. (2017). Activity-Dependent Gating of Parvalbumin Interneuron Function by the Perineuronal Net Protein Brevican. Neuron 95, 639–655.e10.

30. Noutel, J., Hong, Y.K., Leu, B., Kang, E., and Chen, C. (2011). Experience-dependent retinogeniculate synapse remodeling is abnormal in MeCP2-deficient mice. Neuron 70, 35–42.

31. Styr, B., and Slutsky, I. (2018). Imbalance between firing homeostasis and synaptic plasticity drives early-phase Alzheimer’s disease. Nat. Neurosci. 21, 463–473.

32. Tatavarty, V., Torrado Pacheco, A., Groves Kuhnle, C., Lin, H., Koundinya, P., Miska, N.J., Hengen, K.B., Wagner, F.F., Van Hooser, S.D., and Turrigiano, G.G. (2020). Autism-Associated Shank3 Is Essential for Homeostatic Compensation in Rodent V1. Neuron 106, 769–777.e4.

33. Gray, J.M., and Spiegel, I. (2019). Cell-type-specific programs for activity-regulated gene expression. Curr. Opin. Neurobiol. 56, 33–39.

34. Yap, E.-L., and Greenberg, M.E. (2018). Activity-Regulated Transcription: Bridging the Gap between Neural Activity and Behavior. Neuron 100, 330–348.

35. Chen, L.-F., Zhou, A.S., and West, A.E. (2017). Transcribing the connectome: roles for transcription factors and chromatin regulators in activity-dependent synapse development. J. Neurophysiol. 118, 755–770.

36. Ma, H., Khaled, H.G., Wang, X., Mandelberg, N.J., Cohen, S.M., He, X., and Tsien, R.W. (2023). Excitation-transcription coupling, neuronal gene expression and synaptic plasticity. Nat. Rev. Neurosci. 24, 672– 692.

37. Mardinly, A.R., Spiegel, I., Patrizi, A., Centofante, E., Bazinet, J.E., Tzeng, C.P., Mandel-Brehm, C., Harmin, D.A., Adesnik, H., Fagiolini, M., et al. (2016). Sensory experience regulates cortical inhibition by inducing IGF1 in VIP neurons. Nature 531, 371–375.

38. Spiegel, I., Mardinly, A.R., Gabel, H.W., Bazinet, J.E., Couch, C.H., Tzeng, C.P., Harmin, D.A., and Greenberg, M.E. (2014). Npas4 regulates excitatory-inhibitory balance within neural circuits through cell-type-specific gene programs. Cell 157, 1216–1229.

39. Hrvatin, S., Hochbaum, D.R., Nagy, M.A., Cicconet, M., Robertson, K., Cheadle, L., Zilionis, R., Ratner, A., Borges-Monroy, R., Klein, A.M., et al. (2018). Single-cell analysis of experience-dependent transcriptomic states in the mouse visual cortex. Nat. Neurosci. 21, 120–129.

40. Jones, M.W., Errington, M.L., French, P.J., Fine, A., Bliss, T.V., Garel, S., Charnay, P., Bozon, B., Laroche, S., and Davis, S. (2001). A requirement for the immediate early gene Zif268 in the expression of late LTP and long-term memories. Nat. Neurosci. 4, 289–296.

41. Korte, M., Carroll, P., Wolf, E., Brem, G., Thoenen, H., and Bonhoeffer, T. (1995). Hippocampal long-term potentiation is impaired in mice lacking brain-derived neurotrophic factor. Proc. Natl. Acad. Sci. U. S. A. 92, 8856–8860.

42. Plath, N., Ohana, O., Dammermann, B., Errington, M.L., Schmitz, D., Gross, C., Mao, X., Engelsberg, A., Mahlke, C., Welzl, H., et al. (2006). Arc/Arg3.1 is essential for the consolidation of synaptic plasticity and memories. Neuron 52, 437–444.

43. Mikuni, T., Uesaka, N., Okuno, H., Hirai, H., Deisseroth, K., Bito, H., and Kano, M. (2013). Arc/Arg3.1 is a postsynaptic mediator of activity-dependent synapse elimination in the developing cerebellum. Neuron 78, 1024–1035.

44. Yoshihara, S.-I., Takahashi, H., Nishimura, N., Kinoshita, M., Asahina, R., Kitsuki, M., Tatsumi, K., Furukawa-Hibi, Y., Hirai, H., Nagai, T., et al. (2014). Npas4 regulates Mdm2 and thus Dcx in experience-dependent dendritic spine development of newborn olfactory bulb interneurons. Cell Rep. 8, 843–857.

45. Huang, Z.J., Kirkwood, A., Pizzorusso, T., Porciatti, V., Morales, B., Bear, M.F., Maffei, L., and Tonegawa, S. (1999). BDNF regulates the maturation of inhibition and the critical period of plasticity in mouse visual cortex. Cell 98, 739–755.

46. Pelkey, K.A., Barksdale, E., Craig, M.T., Yuan, X., Sukumaran, M., Vargish, G.A., Mitchell, R.M., Wyeth, M.S., Petralia, R.S., Chittajallu, R., et al. (2015). Pentraxins coordinate excitatory synapse maturation and circuit integration of parvalbumin interneurons. Neuron 85, 1257–1272.

47. Inoue, N., Nakao, H., Migishima, R., Hino, T., Matsui, M., Hayashi, F., Nakao, K., Manabe, T., Aiba, A., and Inokuchi, K. (2009). Requirement of the immediate early gene vesl-1S/homer-1a for fear memory formation. Mol. Brain 2, 7.

48. Ramamoorthi, K., Fropf, R., Belfort, G.M., Fitzmaurice, H.L., McKinney, R.M., Neve, R.L., Otto, T., and Lin, Y. (2011). Npas4 regulates a transcriptional program in CA3 required for contextual memory formation. Science 334, 1669–1675.

49. Hashikawa-Hobara, N., Mishima, S., Okujima, C., Shitanishi, Y., and Hashikawa, N. (2021). Npas4 impairs fear memory via phosphorylated HDAC5 induced by CGRP administration in mice. Sci. Rep. 11, 7006.

50. Lin, Y., Bloodgood, B.L., Hauser, J.L., Lapan, A.D., Koon, A.C., Kim, T.-K., Hu, L.S., Malik, A.N., and Greenberg, M.E. (2008). Activity-dependent regulation of inhibitory synapse development by Npas4. Nature 455, 1198–1204.

51. Bloodgood, B.L., Sharma, N., Browne, H.A., Trepman, A.Z., and Greenberg, M.E. (2013). The activity-dependent transcription factor NPAS4 regulates domain-specific inhibition. Nature 503, 121–125.

52. Hong, E.J., McCord, A.E., and Greenberg, M.E. (2008). A biological function for the neuronal activity-dependent component of Bdnf transcription in the development of cortical inhibition. Neuron 60, 610– 624.

53. Chang, M.C., Park, J.M., Pelkey, K.A., Grabenstatter, H.L., Xu, D., Linden, D.J., Sutula, T.P., McBain, C.J., and Worley, P.F. (2010). Narp regulates homeostatic scaling of excitatory synapses on parvalbumin-expressing interneurons. Nat. Neurosci. 13, 1090–1097.

54. Cohen, S.M., Ma, H., Kuchibhotla, K.V., Watson, B.O., Buzsáki, G., Froemke, R.C., and Tsien, R.W. (2016). Excitation-Transcription Coupling in Parvalbumin-Positive Interneurons Employs a Novel CaM Kinase-Dependent Pathway Distinct from Excitatory Neurons. Neuron 90, 292–307.

55. Chew, K.S., Renna, J.M., McNeill, D.S., Fernandez, D.C., Keenan, W.T., Thomsen, M.B., Ecker, J.L., Loevinsohn, G.S., VanDunk, C., Vicarel, D.C., et al. (2017). A subset of ipRGCs regulates both maturation of the circadian clock and segregation of retinogeniculate projections in mice. Elife 6. 10.7554/eLife.22861.

56. Moldavan, M.G., Sollars, P.J., Lasarev, M.R., Allen, C.N., and Pickard, G.E. (2018). Circadian Behavioral Responses to Light and Optic Chiasm-Evoked Glutamatergic EPSCs in the Suprachiasmatic Nucleus of ipRGC Conditional vGlut2 Knock-Out Mice. eNeuro 5. 10.1523/ENEURO.0411-17.2018.

57. Apelblat, D., Roethler, O., Bitan, L., Keren-Shaul, H., and Spiegel, I. (2022). Meso-seq for in-depth transcriptomics in ultra-low amounts of FACS-purified neuronal nuclei. Cell Rep Methods 2, 100259.

58. Liu, S., Cheng, Y., Wang, S., and Liu, H. (2021). Circadian Clock Genes Modulate Immune, Cell Cycle and Apoptosis in the Diagnosis and Prognosis of Pan-Renal Cell Carcinoma. Front Mol Biosci 8, 747629.

59. Hasan, S., van der Veen, D.R., Winsky-Sommerer, R., Hogben, A., Laing, E.E., Koentgen, F., Dijk, D.-J., and Archer, S.N. (2014). A human sleep homeostasis phenotype in mice expressing a primate-specific PER3 variable-number tandem-repeat coding-region polymorphism. FASEB J. 28, 2441–2454.

60. Xue, M., Atallah, B.V., and Scanziani, M. (2014). Equalizing excitation-inhibition ratios across visual cortical neurons. Nature 511, 596–600.

61. Salinas, E., and Sejnowski, T.J. (2000). Impact of correlated synaptic input on output firing rate and variability in simple neuronal models. J. Neurosci. 20, 6193–6209.

62. Chance, F.S., Abbott, L.F., and Reyes, A.D. (2002). Gain modulation from background synaptic input. Neuron 35, 773–782.

63. Cohen-Kashi Malina, K., Tsivourakis, E., Kushinsky, D., Apelblat, D., Shtiglitz, S., Zohar, E., Sokoletsky, M., Tasaka, G.-I., Mizrahi, A., Lampl, I., et al. (2021). NDNF interneurons in layer 1 gain-modulate whole cortical columns according to an animal’s behavioral state. Neuron 109, 2150–2164.e5.

64. Reimer, J., Froudarakis, E., Cadwell, C.R., Yatsenko, D., Denfield, G.H., and Tolias, A.S. (2014). Pupil fluctuations track fast switching of cortical states during quiet wakefulness. Neuron 84, 355–362.

65. Yap, E.-L., Pettit, N.L., Davis, C.P., Nagy, M.A., Harmin, D.A., Golden, E., Dagliyan, O., Lin, C., Rudolph, S., Sharma, N., et al. (2021). Bidirectional perisomatic inhibitory plasticity of a Fos neuronal network. Nature 590, 115–121.

66. Sakata, K., Woo, N.H., Martinowich, K., Greene, J.S., Schloesser, R.J., Shen, L., and Lu, B. (2009). Critical role of promoter IV-driven BDNF transcription in GABAergic transmission and synaptic plasticity in the prefrontal cortex. Proc. Natl. Acad. Sci. U. S. A. 106, 5942–5947.

67. Kohara, K., Yasuda, H., Huang, Y., Adachi, N., Sohya, K., and Tsumoto, T. (2007). A local reduction in cortical GABAergic synapses after a loss of endogenous brain-derived neurotrophic factor, as revealed by single-cell gene knock-out method. J. Neurosci. 27, 7234–7244.

68. Brigidi, G.S., Hayes, M.G.B., Delos Santos, N.P., Hartzell, A.L., Texari, L., Lin, P.-A., Bartlett, A., Ecker, J.R., Benner, C., Heinz, S., et al. (2019). Genomic Decoding of Neuronal Depolarization by Stimulus-Specific NPAS4 Heterodimers. Cell 179, 373–391.e27.

69. Resulaj, A., Ruediger, S., Olsen, S.R., and Scanziani, M. (2018). First spikes in visual cortex enable perceptual discrimination. Elife 7, 1–46.

70. Wen, W., and Turrigiano, G.G. (2024). Keeping Your Brain in Balance: Homeostatic Regulation of Network Function. Annu. Rev. Neurosci. 10.1146/annurev-neuro-092523-110001.

71. Orbán, G., Berkes, P., Fiser, J., and Lengyel, M. (2016). Neural Variability and Sampling-Based Probabilistic Representations in the Visual Cortex. Neuron 92, 530–543.

72. Hengen, K.B., Torrado Pacheco, A., McGregor, J.N., Van Hooser, S.D., and Turrigiano, G.G. (2016). Neuronal Firing Rate Homeostasis Is Inhibited by Sleep and Promoted by Wake. Cell 165, 180–191.

73. Torrado Pacheco, A., Tilden, E.I., Grutzner, S.M., Lane, B.J., Wu, Y., Hengen, K.B., Gjorgjieva, J., and Turrigiano, G.G. (2019). Rapid and active stabilization of visual cortical firing rates across light-dark transitions. Proc. Natl. Acad. Sci. U. S. A. 116, 18068–18077.

74. Dani, V.S., Chang, Q., Maffei, A., Turrigiano, G.G., Jaenisch, R., and Nelson, S.B. (2005). Reduced cortical activity due to a shift in the balance between excitation and inhibition in a mouse model of Rett syndrome. Proc. Natl. Acad. Sci. U. S. A. 102, 12560–12565.

75. Bridi, M.C.D., Zong, F.-J., Min, X., Luo, N., Tran, T., Qiu, J., Severin, D., Zhang, X.-T., Wang, G., Zhu, Z.-J., et al. (2020). Daily Oscillation of the Excitation-Inhibition Balance in Visual Cortical Circuits. Neuron 105, 621–629.e4.

76. Hartzell, A.L., Martyniuk, K.M., Brigidi, G.S., Heinz, D.A., Djaja, N.A., Payne, A., and Bloodgood, B.L. (2018). NPAS4 recruits CCK basket cell synapses and enhances cannabinoid-sensitive inhibition in the mouse hippocampus. Elife 7. 10.7554/eLife.35927.

77. Sharma, N., Pollina, E.A., Nagy, M.A., Yap, E.-L., DiBiase, F.A., Hrvatin, S., Hu, L., Lin, C., and Greenberg, M.E. (2019). ARNT2 Tunes Activity-Dependent Gene Expression through NCoR2-Mediated Repression and NPAS4-Mediated Activation. Neuron 102, 390–406.e9.

78. Padamsey, Z., Katsanevaki, D., Dupuy, N., and Rochefort, N.L. (2022). Neocortex saves energy by reducing coding precision during food scarcity. Neuron 110, 280–296.e10.

79. Pettit, N.L., Yap, E.-L., Greenberg, M.E., and Harvey, C.D. (2022). Fos ensembles encode and shape stable spatial maps in the hippocampus. Nature 609, 327–334.

80. Tanaka, K.Z., He, H., Tomar, A., Niisato, K., Huang, A.J.Y., and McHugh, T.J. (2018). The hippocampal engram maps experience but not place. Science 361, 392–397.

81. Mukherjee, D., Ignatowska-Jankowska, B.M., Itskovits, E., Gonzales, B.J., Turm, H., Izakson, L., Haritan, D., Bleistein, N., Cohen, C., Amit, I., et al. (2018). Salient experiences are represented by unique transcriptional signatures in the mouse brain. Elife 7, 1–20.

82. Whitney, O., Pfenning, A.R., Howard, J.T., Blatti, C.A., Liu, F., Ward, J.M., Wang, R., Audet, J.N., Kellis, M., Mukherjee, S., et al. (2014). Core and region-enriched networks of behaviorally regulated genes and the singing genome. Science 346, 1256780–1256780.

83. Wu, Y.E., Pan, L., Zuo, Y., Li, X., and Hong, W. (2017). Detecting Activated Cell Populations Using Single-Cell RNA-Seq. Neuron 96, 313–329.e6.

84. Ye, L., Allen, W.E., Thompson, K.R., Tian, Q., Hsueh, B., Ramakrishnan, C., Wang, A.-C., Jennings, J.H., Adhikari, A., Halpern, C.H., et al. (2016). Wiring and Molecular Features of Prefrontal Ensembles Representing Distinct Experiences. Cell 165, 1776–1788.

85. Park, E., and Barth, A.L. (2022). IEG expression defines SST neuron ensembles critical for motor learning. Neuron 110, 3222–3224.

86. Hughes, B.W., Siemsen, B.M., Tsvetkov, E., Berto, S., Kumar, J., Cornbrooks, R.G., Akiki, R.M., Cho, J.Y., Carter, J.S., Snyder, K.K., et al. (2023). NPAS4 in the medial prefrontal cortex mediates chronic social defeat stress-induced anhedonia-like behavior and reductions in excitatory synapses. Elife 12. 10.7554/eLife.75631.

87. Yang, J., Serrano, P., Yin, X., Sun, X., Lin, Y., and Chen, S.X. (2022). Functionally distinct NPAS4-expressing somatostatin interneuron ensembles critical for motor skill learning. Neuron 110, 3339–3355.e8.

88. Kyrke-Smith, M., Volk, L.J., Cooke, S.F., Bear, M.F., Huganir, R.L., and Shepherd, J.D. (2021). The Immediate Early Gene Arc Is Not Required for Hippocampal Long-Term Potentiation. J. Neurosci. 41, 4202–4211.

89. Challis, R.C., Ravindra Kumar, S., Chan, K.Y., Challis, C., Beadle, K., Jang, M.J., Kim, H.M., Rajendran, P.S., Tompkins, J.D., Shivkumar, K., et al. (2019). Systemic AAV vectors for widespread and targeted gene delivery in rodents. Nat. Protoc. 14, 379–414.

90. Jaitin, D.A., Kenigsberg, E., Keren-Shaul, H., Elefant, N., Paul, F., Zaretsky, I., Mildner, A., Cohen, N., Jung, S., Tanay, A., et al. (2014). Massively Parallel Single-Cell RNA-Seq for Marker-Free Decomposition of Tissues into Cell Types. Science 343, 776–779.

91. Kohen, R., Barlev, J., Hornung, G., Stelzer, G., Feldmesser, E., Kogan, K., Safran, M., and Leshkowitz, D. (2019). UTAP: User-friendly Transcriptome Analysis Pipeline. BMC Bioinformatics 20, 154.

92. Martin, M. (2011). Cutadapt removes adapter sequences from high-throughput sequencing reads. EMBnet.journal 17, 10–12.

93. Dobin, A., Davis, C.A., Schlesinger, F., Drenkow, J., Zaleski, C., Jha, S., Batut, P., Chaisson, M., and Gingeras, T.R. (2013). STAR: ultrafast universal RNA-seq aligner. Bioinformatics 29, 15–21.

94. Anders, S., Pyl, P.T., and Huber, W. (2015). HTSeq--a Python framework to work with high-throughput sequencing data. Bioinformatics 31, 166–169.

95. Love, M.I., Huber, W., and Anders, S. (2014). Moderated estimation of fold change and dispersion for RNA-seq data with DESeq2. Genome Biol. 15, 550.

96. Pachitariu, M., Stringer, C., Dipoppa, M., Schröder, S., Federico Rossi, L., Dalgleish, H., Carandini, M., and Harris, K.D. (2017). Suite2p: beyond 10,000 neurons with standard two-photon microscopy. bioRxiv, 061507. 10.1101/061507.

97. Scheyltjens, I., Laramée, M.-E., Van den Haute, C., Gijsbers, R., Debyser, Z., Baekelandt, V., Vreysen, S., and Arckens, L. (2015). Evaluation of the expression pattern of rAAV2/1, 2/5, 2/7, 2/8, and 2/9 serotypes with different promoters in the mouse visual cortex. J. Comp. Neurol. 523, 2019–2042.

